# Bipartite viral RNA genome heterodimerization influences genome packaging and virion thermostability

**DOI:** 10.1101/2022.07.29.501896

**Authors:** Yiyang Zhou, Andrew L. Routh

## Abstract

Diverse RNA-RNA interactions occur throughout the lifecycle of RNA viruses, including genome dimerization or multimerization through non-covalent association of two or more genomic segments. Both homo-dimerization of retrovirus genomic RNA and hetero-multimerization of genomic segments of multipartite viruses are recognized as important factors that govern genome packaging. The heterodimer of the bipartite alphanodaviruses represents a unique case as its formation is conserved among different virus species but is only observed within the context of fully assembled virus particles. In spite of this, the RNA sequences involved in heterodimerization are unknown. It also remains unclear whether the formation of this heterodimer might impact any stage of the virus lifecycle. In this study, we used Flock House virus (FHV) as a model system to uncover the nucleotide composition of the heterodimer and dissected the impact of heterodimer formation upon numerous stages of the virus lifecycle. We developed a novel Next-Generation Sequencing (NGS) approach called “*XL-ClickSeq*” to probe candidate heterodimer sequences. We found that the heterodimer is formed via intermolecular base-pairing and its formation is retained in defective virus particles. One heterodimer site was identified by thermodynamic prediction that, upon mutagenic disruption, exhibited significant deficiencies in genome packaging and encapsidation specificity to viral genomic RNA. Furthermore, we demonstrate that disruption of this RNA secondary structure directly impacts the thermostability of mature virus particles. These results demonstrated that the intermolecular RNA-RNA interactions within the encapsidated genome of an RNA virus can have an important impact on virus particle integrity and stability and thus its transmission to a new host.

## Introduction

The packaged genome of RNA viruses encompasses a myriad of both short and long-range RNA-RNA interactions. These RNA-RNA interactions mediate a variety of basic functions in the virus lifecycle including replication, gene expression, and genome packaging (Alvarez Diego, Lodeiro María et al. 2005, Miller and White 2006, Shetty, Stefanovic et al. 2013, Nicholson and White 2014, Newburn and White 2019, Le Sage, Kanarek et al. 2020). An important example includes the non-covalent association of two or more RNA genomic segments in multipartite virus families that are co-packaged into single virus particles. Retroviruses represent the most well-recognized case, in which two copies of the +ssRNA genome are homo-dimerized and encapsidated in virions (Paillart, Marquet et al. 1996, Paillart, Shehu-Xhilaga et al. 2004). In HIV-1, the dimerizing sequences and their protein partners have been well characterized (Moore and Hu 2009). HIV-1 genomic dimerization is crucial for viral assembly and virion maturation, and indispensable for genome reverse transcription and recombination (Paillart, Marquet et al. 1996, Paillart, Shehu-Xhilaga et al. 2004). In a similar fashion, hetero-dimerization occurs between genomic RNA segments of multi-partite viruses such as Bluetongue virus and rotavirus (Krishna and Schneemann 1999, Guenther, Sit et al. 2004, AlShaikhahmed, Leonov et al. 2018, Newburn and White 2019). These transient intermolecular RNA-RNA interactions are required for trans-activation of RNA replication, segment assortment, and genome packaging (Guenther, Sit et al. 2004, AlShaikhahmed, Leonov et al. 2018).

One interesting case of viral genome heterodimerization is presented by *Nodaviridae*: a family of +ssRNA viruses with a small bipartite genome encapsidated into non-enveloped icosahedral particles. Both Flock House virus (FHV) and Nodamura virus, despite their serological and structural differences (Schneemann, Reddy et al. 1998), heterodimerize their bipartite RNA genomes into a single RNA complex upon heating the virions (Krishna and Schneemann 1999). The non-covalent heterodimerized viral RNA can withstand certain denaturing conditions during RNA extraction, as well as native gel-electrophoresis (Krishna and Schneemann 1999). It is unknown whether such a “semi-permanent” genome heterodimerization plays a role in virus life cycles and whether such duplexing is required for the correct encapsidation of each RNA molecule. It is also unknown what sequences contribute to the formation of the heterodimer.

In this study, we use FHV as a model system to reveal the sequences related to heterodimer formation and to demonstrate how such intermolecular RNA-RNA interaction have important roles in multiple stages of the viral lifecycle. FHV is a versatile model system for the study of the structure and molecular biology of non-enveloped viruses. The small bipartite genome encodes the RNA-dependent RNA polymerase (RdRp) on RNA1 (3.1kb) and the structural capsid protein on RNA2 (1.4kb) (Odegard, Banerjee et al. 2010). RNA1 also encodes a subgenomic RNA (RNA3) who’s expression is regulated via a series of cis-acting elements (Lindenbach, Sgro et al. 2002), and yields protein B2 as an essential suppressor of cellular anti-viral silencing mechanisms (Li, Li et al. 2002). The replication of the FHV genome takes place in spherules formed by invaginations of outer mitochondrial membranes (Miller and Ahlquist 2002), where RNA1 replicates independently and trans-activates RNA2 replication (Johnson and Ball 1999, Lindenbach, Sgro et al. 2002). Structurally, the T=3 icosahedral FHV virion packages specifically one molecule of RNA1 and RNA2, whereas the subgenomic RNA3 is excluded (Scotti, Dearing et al. 1983, van de Waterbeemd, Fort et al. 2017). Cryo-EM and X-ray crystallography studies revealed the encapsidated FHV genome forms a highly ordered dodecahedral cage of RNA (Tang, Johnson et al. 2001, Johnson, Tang et al. 2004). It also has been demonstrated that the FHV RNA dodecahedral cage extensively interacts with the capsid shell with a clear structural tropism (Zhou and Routh 2020).

Virus-like particles (VLPs) of FHV can readily be expressed and generated in cell-culture. Interestingly, VLPs predominantly package host RNAs in place of the viral genome (Tihova, Dryden et al. 2004, Routh, Domitrovic et al. 2012, Routh, Domitrovic et al. 2012). In spite of this, the molecular structure of VLPs is indistinguishable from that authentic FHV virions containing the viral genome, and a dodecahedral cage of RNA is also observed. These findings indicate that the structure of mature FHV itself does not prevent the encapsidation of non-viral RNAs and so the virus must employ other strategies to ensure the encapsidation of viral genome. A stem-loop structure on RNA2 has been found to be essential for FHV genome packaging (Zhong, Dasgupta et al. 1992). Additionally, several capsid motifs have been shown to impact RNA recognition (Dong, Natarajan et al. 1998, Schneemann and Marshall 1998, Marshall and Schneemann 2001). Recent studies revealed FHV genome packaging may employ multiple synergetic packaging sites (Zhou and Routh 2020). Despite detailed characterization of the structure of mature FHV particles and the encapsidated genome, the molecular mechanisms that direct the specific and stoichiometric packaging of FHV RNAs still remain elusive.

In this study, we hypothesized that viral genome heterodimerization is formed as a result of RNA1-RNA2 interactions and that these specific intermolecular interactions impact both the efficiency and the specification of genome packaging of the virus. We demonstrate that the heterodimerized RNA can be crosslinked with a psoralen derivative: 4′-aminomethyl trioxsalen hydrochloride (AMT). This suggests the heterodimer is formed as a result of RNA1-RNA2 base pairing. We also introduce a novel Next-Generation Sequencing (NGS) method: ‘*XL-ClickSeq’* (crosslink-ClickSeq), which is designed to specifically probe the double-stranded RNA regions. We found that defective FHV particles retained heterodimer formation, in spite of containing large deletions in each genomic RNA, supporting a biological role of heterodimer.

Thermodynamic prediction revealed one RNA1-RNA2 interaction site (RNA1:1092-1104 and RNA2: 277-288) that contributes to heterodimer formation. Recombinant viruses with disrupted heterodimer base-pairing at this site exhibited significant reduction in packaging efficiency, without compromising the yield of capsid protein or genomic RNAs. Mutant viruses also displayed reduced packaging specificity, which encapsidated a reduced complement of RNA2 and more host RNAs than that of wild-type. We also developed a fluorescence-based method to monitor the thermostability of viral particles. Viruses with disrupted heterodimer base pairs showed significant reduction in thermostability in both molecular and high-throughput assays. These results support an important role for the FHV genome heterodimer in the virus lifecycle, particularly in governing the virus packaging process and structural integrity. The methods and techniques presented in this study shed light on the connection between RNA structure and the virus replication cycles. Importantly, beyond elucidating mechanisms for genome packaging, our findings demonstrate a fundamental and causal link between the higher-order structure of encapsidated genomic RNAs and the structural integrity of virus particles. Therefore, rather than being a passive cargo, the viral genomic nucleic acid itself plays an important role in virus particle integrity and stability and thus its transmission to a new host.

## Results

### FHV heterodimer formation can be thermostabilized by psoralen crosslinking

The stable heterodimerization of each of the two genomic segments of Flock House virus (FHV) to form an RNA1-RNA2 duplex upon heating and cooling of FHV particles has previously been well-described (Krishna and Schneemann 1999). Interestingly, the formation of this heterodimer only occurs within the context of authentic virus particles and is not formed *in vitro* with purified viral RNAs alone (Krishna and Schneemann 1999). This suggests that the assembled virus particles impose one or more well-defined and conserved interaction points between each genomic segment, although the functional importance is not characterized. To address this and to identify the RNA1-RNA2 duplex interaction sites, we recreated the heterodimerization by heating purified FHV virions at 65°C for 10 min, followed by slow cooling at a rate of 1°C/5 sec until 4°C and RNA extraction. Upon electrophoresis on nondenaturing agarose gel, the RNA1-RNA2 heterodimer (hetdim) appeared as the predominant RNA species, with a small amount of monomer RNA1 and RNA2 (**Figure 1a**). The broad molecular weight range of the heterodimer band (3.5k to 5k) indicates the heterodimer may consist of multiple RNA conformations.

**Figure 1.**
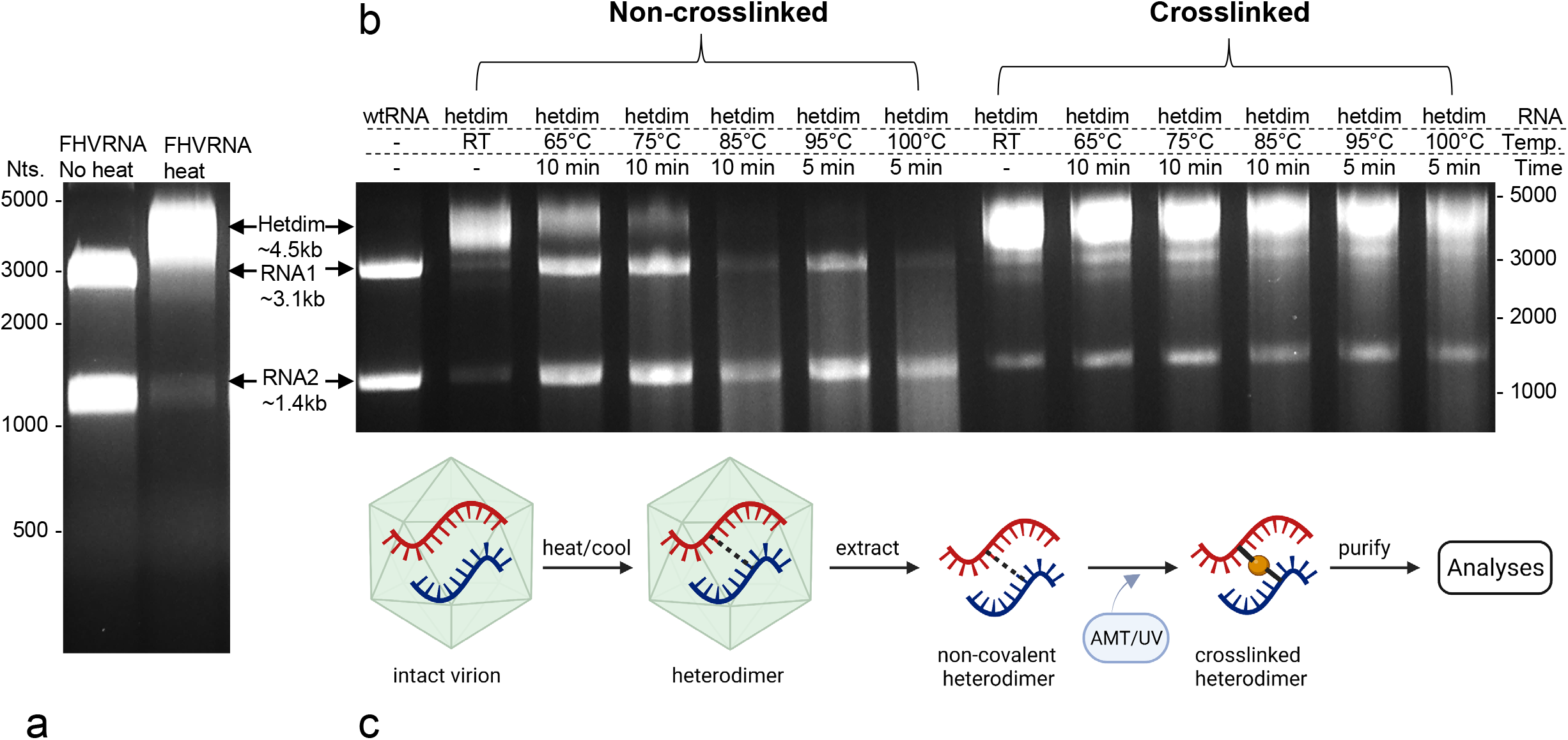
FHV RNA1-RNA2 base pairing contributed to heterodimer formation. **(a)** Upon heating of virions, the majority of FHV RNA1 and RNA2 formed a heterodimerized RNA duplex (hetdim), shown as a band ∼4.5kb on non-denaturing agarose gel. **(b)** Heterodimerized FHV RNA was subjected to 4′-aminomethyltrioxsalen hydrochloride (AMT, psoralen) and UVA light to induce cross-strand pyrimidine crosslinking. The crosslinked heterodimer band can withstand heat up to 100 °C for 5 min, while the non-crosslinked heterodimer bands gradually denatured into separate RNAs with temperature increases. 500ng of total RNA was used in each lane. The molecular weight marker are referenced with ssRNA ladder, which only represent the approximate RNA size estimation under nondenaturing condition. **(c)** Schematic representation of the method for crosslinking FHV heterodimerized RNAs after extraction.

Psoralens are planar photochemical reagents that can intercalate double-stranded nucleic acids and form inter-strand covalent crosslinks between adjacent pyrimidines upon UVA irradiation (Cimino, Gamper et al. 1985). To investigate whether FHV RNA heterodimer is formed as a result of RNA1-RNA2 base pairing, a psoralen derivative 4′-aminomethyl trioxsalen hydrochloride (AMT) was supplemented to extracted RNAs from heated virions, which consists mainly of heterodimerized viral RNA (**Figure 1b**). After crosslinking with 365nm UVA light, the AMT-crosslinked heterodimer became resistant to heat denaturation at temperatures up to 100°C (**Figure 1b**). In comparison, non-crosslinked heterodimer began to denature at 65°C and were completely monomerized at 85°C. This demonstrates that the heterodimerization is a result of inter-strand base-pairing between RNA1 and RNA2.

### ‘*XL-ClickSeq*’ to identify dsRNA interaction sites

Having confirmed that AMT-crosslinking preserves the interaction between RNA1 and RNA2 in the FHV heterodimer, we employed Next-Generation Sequencing (NGS) approaches to identify the interaction site(s). We developed a novel approach that we call “*XL-ClickSeq*”, which combines psoralen crosslinking (“XL”) and “ClickSeq” (Routh, Head et al. 2015) to identify the sequence of AMT-crosslinked double-stranded structures of RNA (**Figure 2a**). Random-primed ClickSeq relies on the stochastic incorporation of 3’-azido-deoxynucleotides (AzNTPs) to randomly terminate cDNA synthesis during reverse-transcription (RT) of unfragmented RNA templates and subsequently ‘click-ligates’ (Kolb, Finn et al. 2001) a 5’-alkyne-functionalized oligonucleotide containing the Illumina sequencing adaptor. In XL-ClickSeq, unfragmented RNA template was first AMT-crosslinked. The AMT-crosslink induced cross-strand covalent bonds which prevents the elongation of the nascent cDNA past the crosslink and therefore prevents the required incorporation of an AzNTPs near this site. This prevents click-ligation to the click-adapter and thus prevents PCR amplification and subsequent sequencing on an Illumina flowcell. This results in a loss of sequence read coverage at the 3’ of crosslinking sites. By contrasting to the normal coverage of non-crosslinked genomic regions, steep drops in genomic coverage indicate crosslinked dsRNA regions at near-nucleotide resolution.

**Figure 2.**
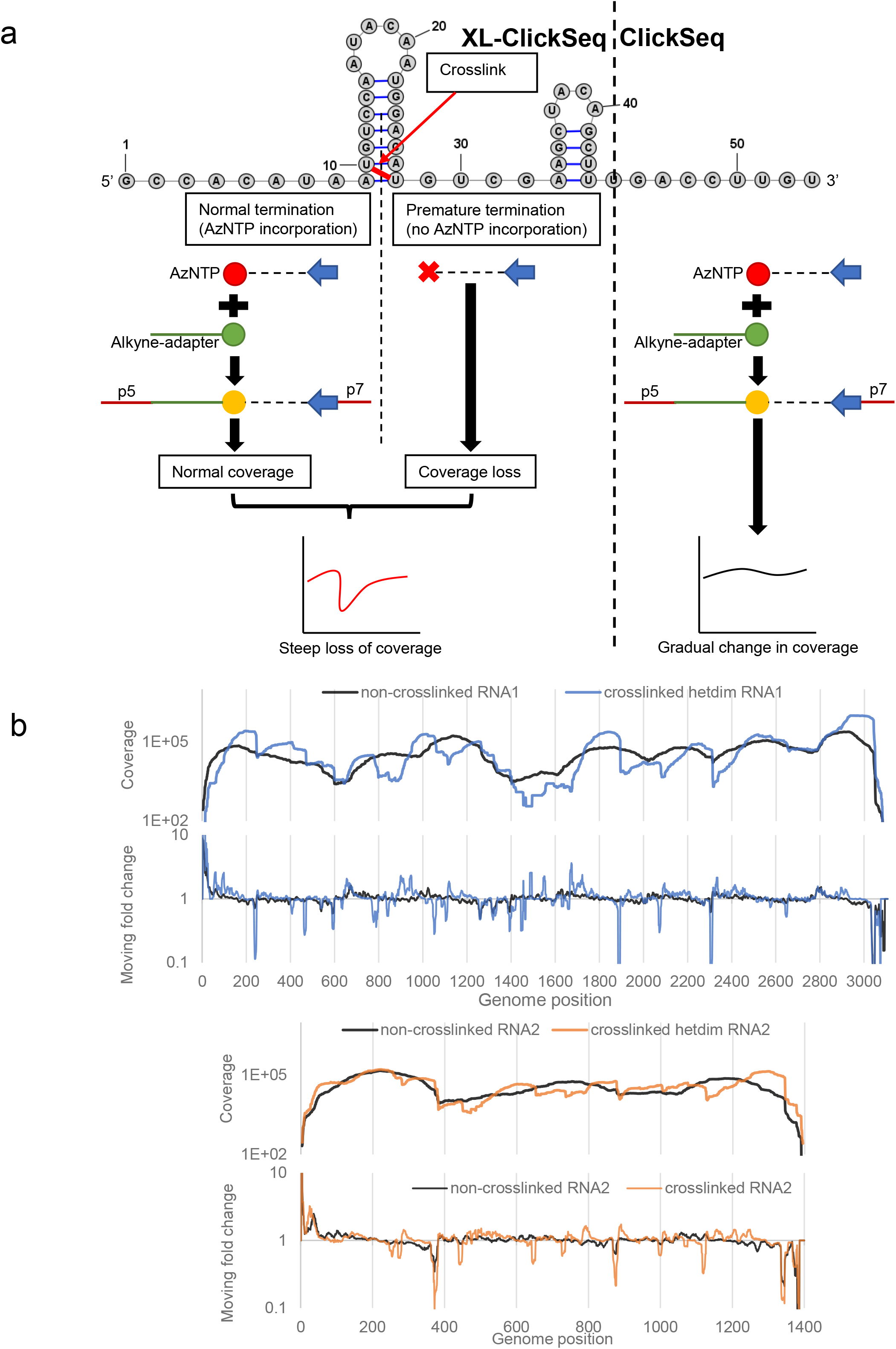
“XL-ClickSeq” revealed heterodimer candidate sites. **(a)** “XL-ClickSeq” method combines psoralen crosslinking (“XL”) with “ClickSeq” to specifically probe crosslinked double-stranded RNA regions. RNA molecule is UV-crosslinked with psoralens (AMT). The cross-stranded covalent bonds prevent AzNTP incorporation during RT, which subsequently impedes the click-ligation to an aklyne-functionalized Illumina adapter. The RNA base pairing is thereafter revealed by the sharp coverage loss near crosslinking sites. **(b)** XL-ClickSeq of heterodimerized FHV RNA revealed several sites with sudden loss of coverage, indicating double-stranded RNA duplexes.

Heterodimerized FHV RNAs were extracted from heated virion and AMT-crosslinked (**Figure 1b**). XL-ClickSeq of crosslinked heterodimer revealed that numerous genomic locations showed apparent loss of coverage, which is distinctively absent from non-crosslinked control RNAs (**Figure 2b**). This is further highlighted by moving fold change (10nt interval, **Figure 2b**). Sharp changes in coverage are characteristic of XL-ClickSeq and indicate the location of crosslinked dsRNA sites. With the XL-ClickSeq data of heterodimer, we applied a manual threshold to distinguish dsRNA sites from background: a candidate site needs to contain consecutive (greater than 6) nucleotides that have a moving fold change reduction (10nt. interval) greater than 25% (**Figure S1**). We also excluded the sites located within 3’-most 100nts of both RNAs, since reduced genomic coverage at the 3’ termini is common in RNAseq (Routh, Head et al. 2015) and such loss of coverage is seen on both crosslinked RNA or non-crosslinked control. We found a total of 18 crosslinking sites on RNA1 and 9 on RNA2, respectively. It is important to note that the revealed dsRNA sites do not differentiate between intramolecular or intermolecular base-pairing.

### FHV heterodimer formation is conserved in defective particles

We have previously characterized the emergence and evolution of defective viral genomes of FHV during serial passaging in cell-culture (Jaworski and Routh 2017). The “mature” defective RNAs (D-RNAs) of FHV are characterized by large deletions of genomic RNA but retain important functional RNA motifs such as those required for RNA replication and encapsidation. To determine whether the RNA1-RNA2 intermolecular interactions are also conserved in ‘mature’ D-RNAs, we investigated whether FHV heterodimer can be formed in defective particles (defective particles are defined as assembled virions that encapsidate D-RNAs). Stocks of defective FHV particles were obtained from the supernatant from drosophila S2 cells in culture after 7^th^ serial rounds of passaging (P7). Nanopore sequencing revealed that P7 FHV virus stock consisted of 92% D-RNA1 and 88% D-RNA2 (**Figure S2**). These D-RNAs are characterized by four predominant genomic deletions (nt. 318-940 and nt. 1261-2265 on RNA1, nt. 251-512 and nt. 717-1217 on RNA2) (**Figure S2**). In comparison, early passage of FHV (e.g. P2) comprised of less than 7% of defective RNA1 and 4% defective RNA2 (Jaworski and Routh 2017), and hence considered wild type (wt) genome in this study.

Surprisingly, despite the truncated genome, the P7 defective FHV particles were still able to form heterodimer upon heating (**Figure 3a**). On non-denaturing agarose gel, the predominant P7 heterodimer species migrated similarly to that of wild type (P2) at 4.5kb, although there could be uncharacterized P7 heterodimer species at lower molecular weight. This suggests that in defective FHV particles, the heterodimer may comprise two copies of truncated RNA1 and RNA2 molecules, since the truncated D-RNA1 is approximately half the molecular weight of full length.

**Figure 3.**
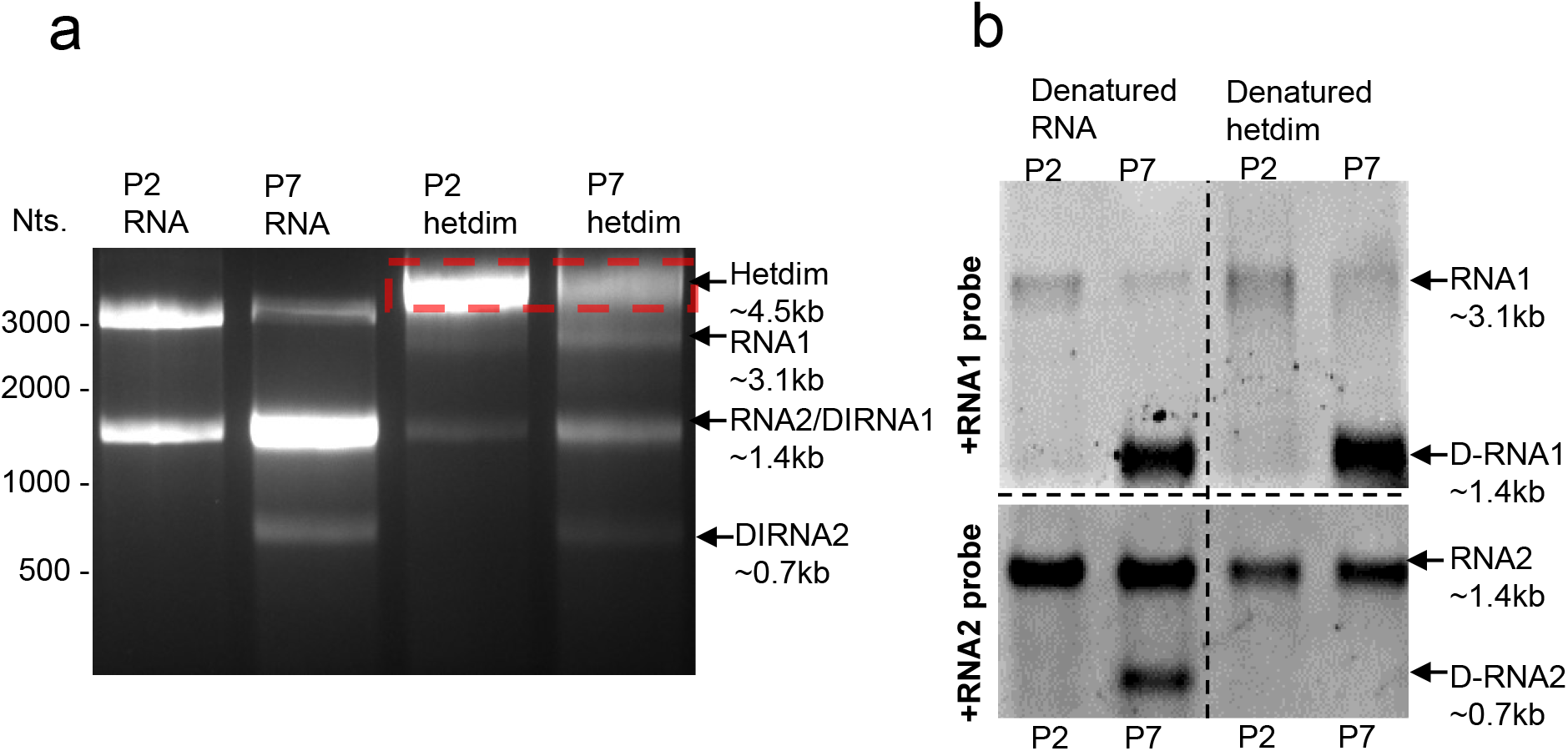
FHV RNA heterodimerization is conserved in defective particles. **(a)** Both P2 and P7 FHV viruses can form heterodimer, despite P7 virus comprising primarily of defective FHV particles, in which RNA genomes contain large deletions (D-RNA1 and D-RNA2). The heterodimer bands (red box) were excised and RNAs were recovered and denatured for subsequent northern blot assay. **(b)** Northern blots of non-heterodimerized RNA (denatured RNA) and heterodimerized RNA (denatured hetdim) from P2 and P7 particles. The D-RNAs from P7 particles are clearly shown. Only D-RNA1 can be detected from P7 particle heterodimer band. The molecular weight marker is referenced with ssRNA ladder, which only represent the approximate RNA size estimation under nondenaturing condition.

To confirm the heterodimer indeed consists of both RNA1 and RNA2, the predominant heterodimer species of both wild type (P2) and defective (P7) viruses were extracted from non-denaturing agarose gel (red box in **Figure 3a**). The retained heterodimer RNAs were then denatured and analyzed by northern blot using probes against +ssRNA1 and +ssRNA2 (**Figure 3b**). D-RNA1 (∼1.4kb) and D-RNA2 (∼0.7kb) are readily detectable in P7 D-FHV. Surprisingly, the P7 heterodimer consisted of full length RNA1, full length RNA2, D-RNA1, but not D-RNA2. The exclusion of D-RNA2 from P7 heterodimer suggests that the heterodimerization may be interfered by the deletions in D-RNA2, while unaffected by the deletions in D-RNA1.

### Potential heterodimer sites and disruptive mutants

The previous experiments provide a range of constraints on where in the FHV genome the RNA1-RNA2 duplex interaction sites must occur. The presence of FHV D-RNA1 and absence of D-RNA2 in P7 heterodimers (**Figure 3b**) indicate that at least one heterodimer site in RNA1 resides in the conserved regions of D-RNA1 (RNA1 nts. 1-317, 941-1260, 2266-3107), while the other complementary site in RNA2 must reside in a region deleted in D-RNA2 (RNA2 nts. 251-512, 737-1217). Therefore, of the 27 crosslinking sites identified by “XL-ClickSeq”, we further refined our selection to 6 sites on RNA1 and 6 sites on RNA2, that are found in these regions and hence may contribute to the formation of RNA1-RNA2 heterodimer (**Figure S3**).

In order to understand which sites can form intermolecular base-pairing duplex(es), we used the “*bifold*” function of *RNAstructure* (Reuter and Mathews 2010) to perform thermodynamic prediction of these candidate sites for both intermolecular and intramolecular base-pairing probabilities (**Figure S3**). Each RNA1 candidate site as well as the flanking sequence (71-120nts. in length) was cross-matched with each RNA2 candidate site and flanking sequence to investigate whether the selected candidate segments have greater potential to form intermolecular duplexes (RNA1-RNA2) than intramolecular ones (RNA1-RNA1 or RNA2-RNA2), whether the formed intermolecular base-pairs are at the predicted crosslinking sites, and whether these base pairs exhibit cross-linkable pyrimidines (**Figure S3**). One particular match (RNA1 nt.1102 and RNA2 nt.280) resulted in a 9 base-pair RNA duplex between RNA1:1094-1102 and RNA2:277-285, which is located at the proximity of predicted crosslinking site **(Figure 4a)**. This double-stranded stem is comprised of 4 pairs of cross-stranded U-U and U-C pairs that are available for AMT-crosslinking (Garrett-Wheeler, Lockard et al. 1984, Cimino, Gamper et al. 1985). Beyond the 9 base-pairing RNA stem, RNA1-RNA2 duplexing is also predicted in adjacent sequences. Between RNA1: 1075-1118 and RNA2: 261-307, 66% of the bases are predicted to be involved in RNA1-RNA2 base-pairing.

**Figure 4.**
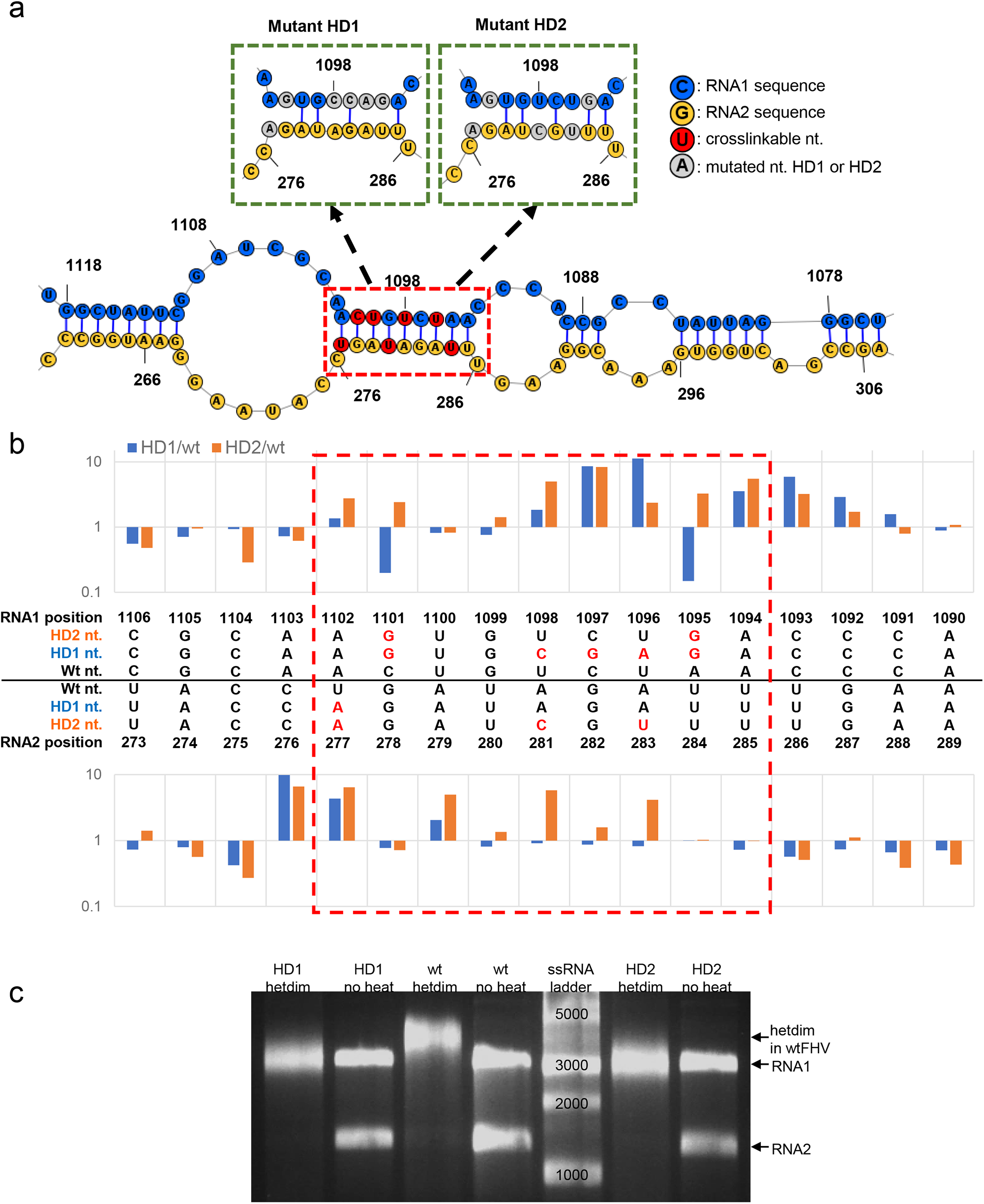
Potential heterodimer sites and disruptive mutants. **(a)** The predicted RNA1: 1093-1104 and RNA2: 276-286 base-pairing (red box, based on cross-matching results from **Supplemental Figure 3**) contains several cross-strand pyrimidines available for AMT crosslinking (U-U and C-U). Two mutant viruses were created (HD1 and HD2), with synonymous mutations designed to disrupt the predicted base pairing in this region. (b) DMS-MaPseq of mutated viruses demonstrated more unpaired bases in the heterodimer stem (red box) as well as flanking sequences. (c) Upon heating, mutant (HD1 and HD2) viral RNAs can still be heterodimerized, but with altered conformation, which migrated faster than heterodimer from wt virus in a native agarose gel.

To scrutinize this potential dimerization site between RNA1:1094-1102 and RNA2:277-285, we designed two mutant viruses (HD1 and HD2) that introduced synonymous mutations to disrupt the predicted base-pairing of this double stranded intermolecular stem. HD1 contained 5 nts. substitutions on RNA1 and 1 nt substitution on RNA2, to predictively disrupt 5 base pairs. HD2 contained 2 and 3 nts. substitutions on RNA1 and RNA2, respectively, to predictively disrupt 4 base pairs (**Figure 4a**). After transfection, the mutant viruses were continuously passaged. Illumina sequencing confirmed that the mutations were preserved in passage 2 (P2) viruses (**Figure S4**).

We first sought to confirm whether RNA secondary structure in the predicted heterodimer region was disrupted by the synonymous mutations in the HD1 and HD2 virus genomes. WT, HD1 and HD2 viruses were purified with polyethylene glycol (PEG) precipitation, sucrose gradient and molecular weight filteration (as previously described (Zhou and Routh 2020)), and then subjected to in-virion dimethyl sulfate mutational profiling with sequencing (DMS-MaPseq) (Umeyama and Ito 2017, Zhou and Routh 2020). We compared the DMS-MaPseq signal between mutant and wild-type viruses at the predicted heterodimer dsRNA regions (9bps, RNA1: 1094-1102; RNA2:277-285, red box of **Figure 4b)**. 7 nucleotides in HD1 and 14 nucleotides in HD2 showed increased DMS signal (defined here as the nucleotide error rate) relative to wild type virus. Furthermore, in both mutants, additional nucleotides also exhibited higher DMS signal at the 5’ proximal of the predicted heterodimer stem. This indicates the introduced mutations disrupted the base-pairing of the 9 bps heterodimer stem as well as some flanking bases. It is possible that the disrupted RNA base pairs in these mutants underwent compensatory conformational changes to reform base-pairing with other RNA regions. This may explain why HD1 had fewer single-stranded bases than HD2, despite bearing more disruptive mutations.

The conformational change of HD1 and HD2 could also be observed when these mutant viruses were heated to extract heterodimer (**Figure 4c**). Despite disruption of the identified heterodimer stem, both HD1 and HD2 still formed heterodimers. This indicates there are multiple heterodimerization sites beyond the predicted region. Importantly, the heterodimers of both HD1 and HD2 mutant migrated faster than that of wt virus on non-denaturing agarose gel (**Figure 4c**). This suggests the disrupted stem of HD1 and HD2 triggered significant RNA conformational changes, which led to the observed electrophoresis mobility shift. Previously, it has been demonstrated that local RNA substitutions can lead to significant reconfiguration of the genomic RNA structure (Zhou and Routh 2020, Wang, Sotcheff et al. 2022). The heterodimer bands of HD1 and HD2 were extracted, denatured, and northern blot analyzed to verify that they indeed consisted of both RNA1 and RNA2 (**Figure S5**).

As stated above, it is conceivable that there are multiple interaction sites between RNA1 and RNA2, in addition to the discovered stem (RNA1: 1093-1104 and RNA2: 276-286). To confirm this, we designed a molecular approach using DNA-oligo-guided site-specific cleavage by RNase H (**Figure S6**). We designed DNA probes to specific sites flanking RNA1: 1093-1104 and confirmed their site-specific cleavage of monomerized viral RNAs (**Figure S6**). However, the heterodimerized viral RNA resisted perturbation, as both RNA1 and RNA2 still migrated as a single heterodimer band in the native agarose gel. This suggests that there are additional undiscovered RNA1-RNA2 interaction sites.

### Disrupted heterodimer impedes virion assembly and genome packaging specificity

We further sought to characterize HD1 and HD2 mutants to understand if the disrupted heterodimer site could impact virus lifecycle and fitness. AlamarBlue assays were conducted to compare the cytotoxicity of mutant viruses to wild type. After transfecting equal amounts of plasmids containing sequences for wtFHV, HD1 and HD2, the cells transfected by HD1 and HD2 mutants (P0) showed significantly reduced cytotoxicity than that of wild type transfectant (**Figure 5a**), indicating both HD1 and HD2 might propagate slower than wt. Naïve cells were inoculated with equal volume of transfected cell culture (P0). The resultant P1 culture was analyzed with western blot to investigate the capsid yield in cytosol and supernatant (**Figure 5b, c**). Interestingly, neither mutant resulted in significantly different yield of capsid in cytosol (**Figure 5b**). In contrast, both HD1 and HD2 yielded significantly reduced capsid in supernatant fraction (**Figure 5c, Figure S7**), where mature virus accumulates. The reduced supernatant capsid yields are consistent with reduced cytotoxicity of mutants (**Figure 5a**). The reduced virion production of HD1 and HD2 was also confirmed by quantitating the yield of P2 virions after sucrose gradient purification, which was diminished for both the HD1 and HD2 mutants (**Figure 5d**). Overall, this suggests that HD1 and HD2 mutations results in a deficiency in assembling mature virions that escape infected cells, despite HD1 and HD2 expressing comparable amounts of capsid proteins intracellularly to wt.

**Figure 5.**
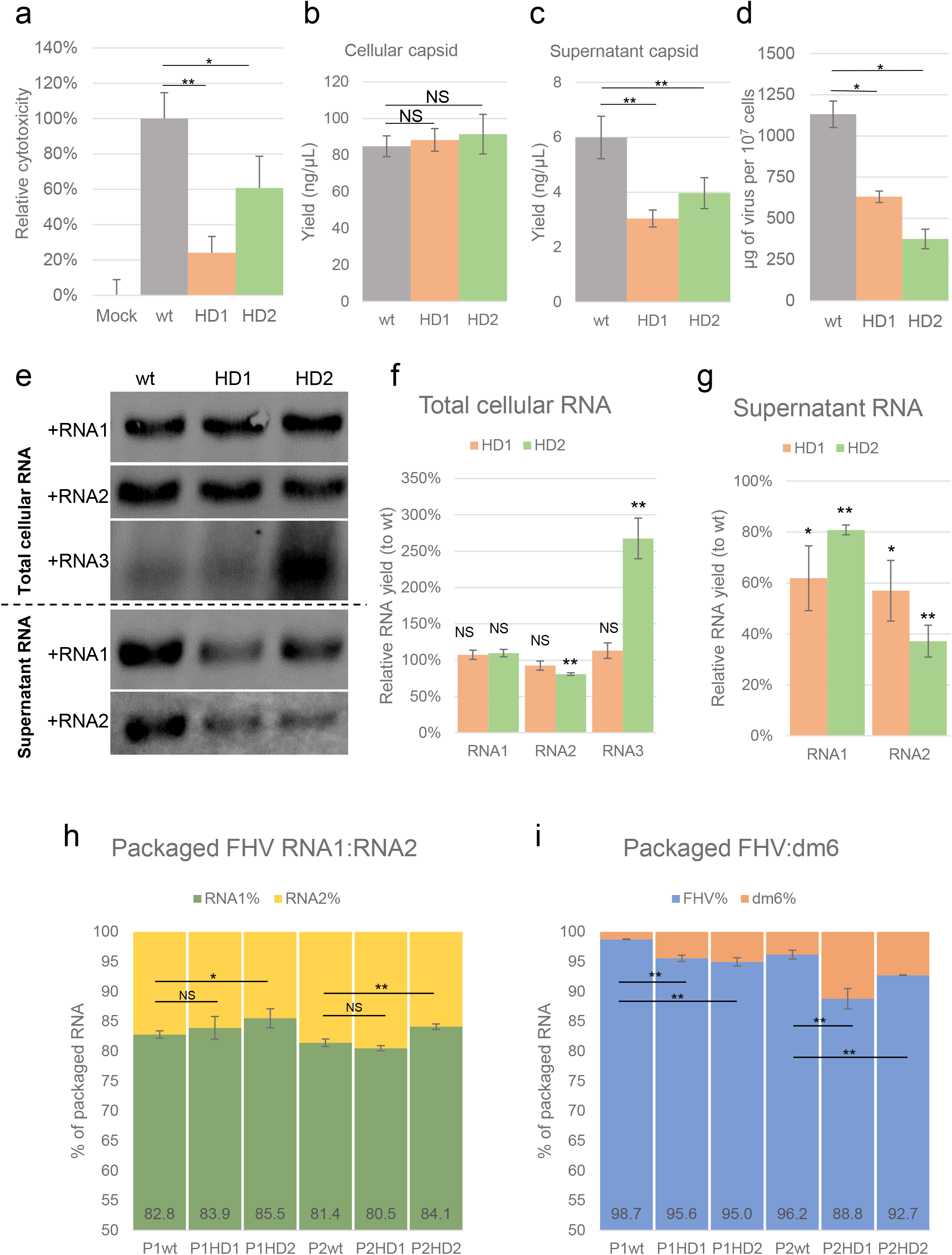
Disrupted heterodimer impedes virion assembly and packaging specificity. **(a)** alamarBlue assay revealed that both mutant viruses (HD1 and HD2) showed significantly reduced cytotoxicity than that of wild type. **(b)** Comparable amounts of capsid were detected by western blot assays in cells infected by wt or mutant viruses. **(c)** Mutant viruses yielded significantly reduced capsid in supernatant. **(d)** Both mutant viruses also showed reduced yield in mature virions. **(e)** Northern blot of the infected cells or supernatant using probes against RNA1 or RNA2. **(f-g)** HD2 mutant virus triggered a significant increase in sgRNA3 abundance in infected cells, which may be the cause of the slight but significant reduction of RNA2. **(h)** HD2 mutant encapsidated significantly more RNA1 and less RNA2 than that of wild type in both P1 and P2 viruses. **(i)** Both HD1 and HD2 packaged significantly more host RNAs than that of wt. (NS: not significant; *: p<0.05; **: p<0.01)

We next sought to determine whether the mutated heterodimer stem would impact viral replication and RNA packaging. 1 μg of total cellular RNA or 300 ng of supernatant RNA were analyzed by northern blot with ssRNA probes specific for each viral RNA (**Figure 5e**). Interestingly, HD2 accumulated significantly more sub-genomic RNA3 in cells and a slight but significant reduction of RNA2 (**Figure 5e, f**). The negative yield correlation between RNA2 and subgenomic RNA3 is consistent with previous findings in FHV and other alphanodaviruses, which report that the replication of RNA2 not only is trans-activated by sgRNA3, but also suppresses the accumulation of sgRNA3 after being trans-activated (Guarino, Ghosh et al. 1984, Zhong and Rueckert 1993, Eckerle and Ball 2002, Albariño, Eckerle et al. 2003, Eckerle, Albarino et al. 2003). This suggests the disrupted heterodimer stem (RNA1:1094-1102 and RNA2: 227-285) may involve a novel function in the coordination of RNA2 and sgRNA3 replication. In supernatant fractions, both HD1 and HD2 showed significantly reduced RNA1 and RNA2 accumulation than that of wt (**Figure 5e, g**), consistent with the lower yield of mature virions from mutant viruses.

We then investigated whether the disrupted heterodimer stem would impact the virus genome packaging. P1 and P2 viruses of HD1, HD2, and wt viruses were collected and quintuple-purified to ensure minimal non-viral RNA content. The encapsidated RNAs were extracted from P1 and P2 viruses and characterized by random-primed RNAseq to determine the relative abundance of encapsidated virus and host RNAs (Routh, Domitrovic et al. 2012, Routh, Domitrovic et al. 2012). We observed that HD2 packaged significantly more RNA1 and less RNA2 than that of wt in both P1 viruses and P2 viruses (**Figure 5h**). This is consistent with the reduced cellular RNA2 yield associated with HD2 (**Figure 5e**). In mutant viruses, both HD1 and HD2 showed significantly increased host RNAs encapsidation in both P1 and P2 viruses (**Figure 5f**). Taken together, this suggests the heterodimer stem plays a role in ensuring FHV genome packaging specificity.

### Disrupted heterodimer reduced virion thermostability

We next sought to understand whether disruption of heterodimer stem could interfere with virion thermostability. We designed a molecular assay (**Figure 6a**), in which 50 μg of the quintuple-purified virions were heated at various temperatures (from 45 °C to 95 °C) for 15 mins and subsequently loaded to a 1% non-denaturing agarose gel pre-stained with nucleic acid dye. After imaging, the same gel was post-stained with Coomassie blue to reveal protein content. We expected that intact virions would be less permeable to nucleic acid dye and preclude fluorescent staining, whereas heat-treated particles would lose their structural integrity and thus be permeable to the intercalating dye resulting in an increase in measured fluorescence. We observed that both HD mutants were more permeable than wt to nucleic acid stain even at RT, which is characterized by brighter fluorescence of the RNA-capsid complex band(s). Such permeability difference was consistent when heating particles up to 75 °C. The most substantial difference could be observed at 55 °C, at which temperature both HD1 and HD2 exhibited higher nucleic acid dye permeability, greater amount of RNA-capsid complex, greater amount of fully disassociated proteins (protein-stained gel) and fewer intact particles (protein-stained gel). All particles appeared to lose integrity at 65 °C, at which point fluorescence peaked. However, at 75 °C, the RNA-capsid complexes of wt, HD1, HD2 showed different electrophoretic mobility shift patterns, indicating different degrees of protein-nucleic acid separation. Above 85°C, all viruses further disintegrated, and no difference was distinguishable between mutant(s) and wt. Despite the high temperature, there was still evidence for retained RNA-capsid interactions as multiple RNA species were observed to migrate slower than purified viral RNA1 or RNA2. It has been previously demonstrated that FHV virion is comprised of extensive RNA-capsid interactions (Zhou and Routh 2020) that may contribute to the heat-resistance observed in this study.

**Figure 6.**
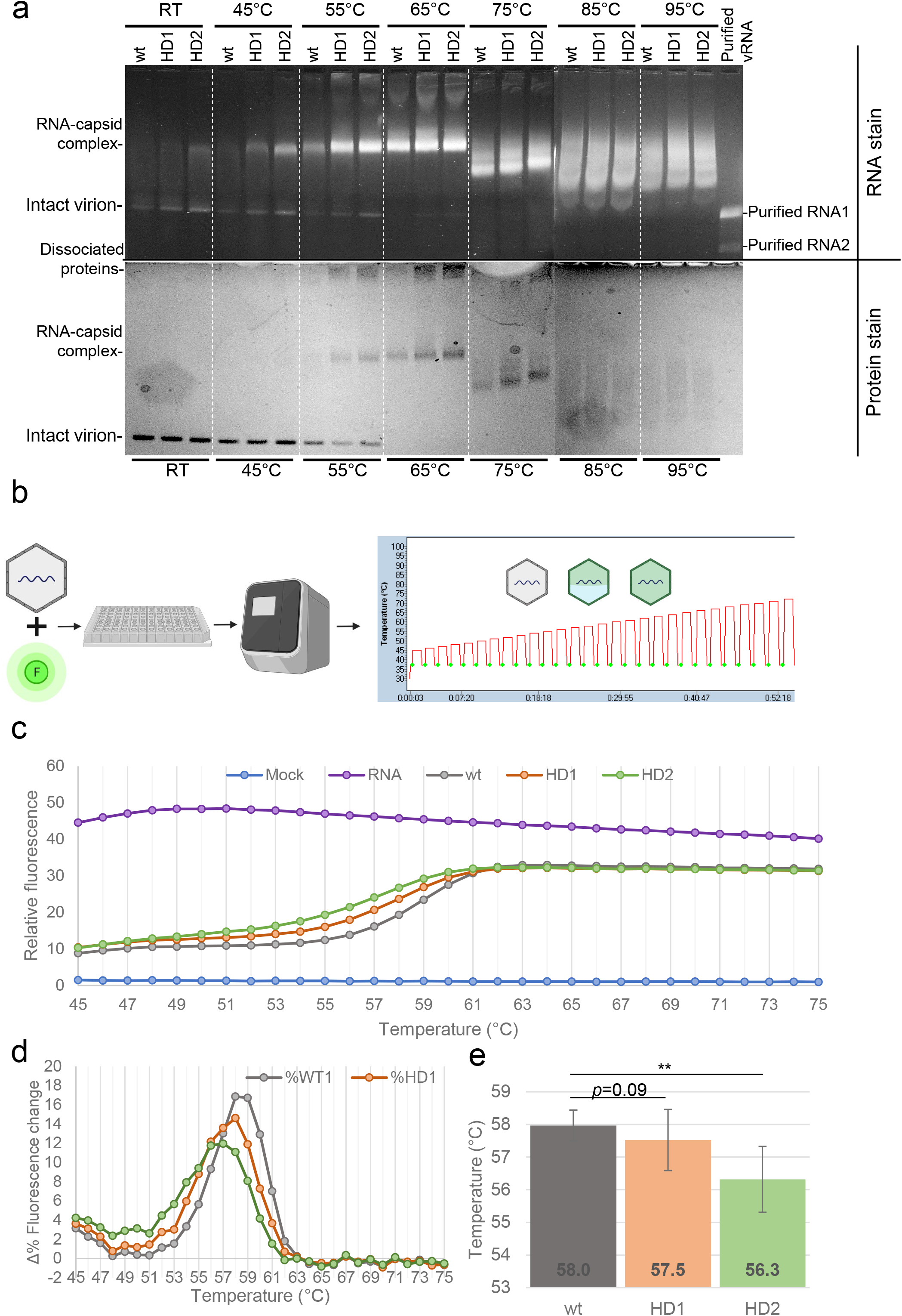
HD1 and HD2 mutants exhibit reduced virion thermostability. (**a**) Wt, HD1 and HD2 particles were heated for 15 min at the temperatures indicated and loaded onto a non-denaturing agarose gel containing nucleic acid stain, which was subsequently post-stained with Coomassie blue to reveal protein content. HD1 and HD2 constantly showed greater permeability of nucleic acid stain up to 75°C. At 55°C, HD1 and HD2 showed substantially reduced amount of intact virion and increased amount of RNA-capsid complex. At 75°C, RNA-capsid complex of three particles showed different mobility shift patterns, indicating further dissociation between protein and RNA. No difference among wt, HD1 and HD2 particles were observed above 85°C. (**b**) A high-throughput fluorimetry assay was designed to monitor thermostability changes of virus particles. Purified mutant or wt virions were mixed with a fluorescent nucleic acid dye and heated for 1 min per Δ1°C, ranging from 45°C to 95°C. Each heating cycle was followed by a rapid cooling to 37°C and a brief incubation (10 s) at 37°C. Green dots represent measurement of relative fluorescence, which is at the end of each colling cycle at 37°C. (**c**) In comparison to wtFHV, both HD1 and HD2 showed increased relative fluorescence at lower temperatures. (**d**) Δ%fluorescence/°C demonstrated that with wtFHV, fluorescence increase started at 54°C and peaked at 60°C. HD1: fluorescence increase started at 52°C and peaked at 59°C.HD2: fluorescence increase started at 49°C and peaked at 57°C. (**e**) Temperature required to reach 50% of relative fluorescence increase.

To test virion thermostability in a high throughput manner, we developed a fluorometric method on a qRT-PCR machine to detect fluorescence increase during particle disassembly upon temperature increases (**Figure 6b**). Similar to the gel electrophoretic approach above, this measures fluorescence change after the heat-induced permeability of virus particles to intercalating nucleic acid dyes. Following each heating step (1 min), relative fluorescence was measured after a rapid cooling to 37 °C (10 s) to remove temperature-dependent effects of dye fluorescence. With this method, we monitored the thermostability changes of wt, HD1 and HD2. The fluorescence increases of all three particles showed a sigmoidal curve (**Figure 6c**). For all three particles, we observed a rapid increase in fluorescence at approximately 51 °C, and a plateau of fluorescence above 61 °C **(Figure 6c, d)**. For wt virions, the greatest rate of fluorescence change was observed at 58-59 °C, indicating the temperature that leads to greatest disassembly of the virus particle and exposure of genomic content to the intercalating dye (**Figure 6d**). This is consistent with our native gel electrophoresis assay that also indicated particle disassembly at between 55 and 65 °C (**Figure 6a**). Interestingly, both HD1 and HD2 showed an increase in fluorescence at lower temperatures than wt (**Figure 6c, d**) and consistently showed more fluorescence than wt between 45-63 °C. Above 63 °C, the relative fluorescence of all three particles plateaued and were equal. This also indicates equimolar pooling of particles. By comparing the Δ rates of particle disassembly (defined here as the Δ percentage of relative fluorescence change, **Figure 6d**), HD1 exhibited the greatest rate of fluorescence increase at 58°C, which rapidly drops thereafter. HD2 showed the greatest rate of fluorescence increase at 57 °C. The melting temperature of a particle (Tm) is defined in this study as the temperature required to reach 50% fluorescence increase of the particle (**Figure 6e**). On average, both HD1 and HD2 showed lower Tm(s) than that of wt. HD2 showed a significantly reduced Tm of 56.3 °C, suggesting that the disrupted heterodimer stem in HD2 significantly compromised the thermostability of the virion.

Altogether, these results demonstrate differences in thermostability between wt and the mutant HD1 or HD2 particles as the result only of synonymous changes in the genomic RNA sequences that disrupt RNA higher order structure inside the virus particle in the absence of any change in the virus capsid shell itself.

## Discussion

### FHV heterodimer and biological importance

Prior to this study, a biological role for the nodavirus genome heterodimer had only limited evidential support (Krishna and Schneemann 1999), including: (1) that the formation of RNA heterodimer is conserved among different viruses of the nodavirus family, such as Nodamura virus and Flock House virus; and (2) that the formation of FHV RNA heterodimer necessitated virion structure. The latter point is in clear contrast to the genome homo-dimerization of retroviruses, in which two identical genomic RNA can dimerize *in vitro* spontaneously (Feng, Copeland et al. 1996). It is possible that a specific virion architecture or particular capsid-RNA association is required for heterodimer formation.

In this study, we provide additional evidence to support a biological role for the FHV heterodimer. We found that: (1) heterodimer is formed as a result of RNA1-RNA2 intermolecular base-pairing and can be crosslinked with AMT (**Figure 1b**); (2) the heterodimer formation is conserved in defective virus particles, despite large internal deletions in the defective genomes (**Figure 3**); (3) the disruption of one heterodimer stem resulted in significant conformational change of the heterodimer (**Figure 4c**); (4) such disruption further attenuated virion assembly (**Figure 5a-d**), RNA replication (**Figure 5e-g**), genome packaging specificity (**Figure 5h-i**) as well as the thermostability of virion (**Figure 6**). Altogether, these observations demonstrate that the intermolecular base-pairing of genomic RNAs is essential to the virus lifecycle, and that disrupting such interactions impacts virus fitness.

To study and observe the heterodimer, we found that specific the RNA extraction method and conditions were critical. Following the heating and slow cooling of virion, a number of commercially available RNA extraction/purification kits/methods were tested. In this study, heterodimer could only be recovered with two methods: (1) proteinase K digestion followed by solid phase reversible immobilization (SPRI) bead pull-down and ethanol wash (reduced yield); and (2) a silica-based RNA purification kit (Quick-RNA Viral kit, Zymo Research), which uses sodium iodide as the leading chaotic reagent (used throughout this study). Interestingly, other silica-based RNA extraction kits (e.g. RNA Clean & Concentrator and Direct-zol RNA prep from Zymo Research) or Trizol-based (Invitrogen) phase separation methods failed to maintain the RNA1-RNA2 association.

It is conceivable that FHV genome consists of multiple heterodimer sites beyond those identified in this study. The disruption (**Figure 4c**) or RNase H digestion (**Figure S6**) both sides of a single heterodimer site was demonstrated to be insufficient to fully disassociate RNA molecules. The existence of numerous heterodimer sites and the extensive RNA1-RNA2 interactions is also supported by previous findings that the majority of packaged FHV RNAs were in double-stranded conformation (Zhou and Routh 2020) and that the FHV genome forms a highly ordered dodecahedral RNA cage inside virion (Tang, Johnson et al. 2001).

We designed synonymous mutations to disrupt the predicted base pairing at the heterodimer stem (RNA1: 1094-1102; RNA2:277-285). Such disruption was determined to have little impact upon the replication of viral RNAs and subsequent expression of viral capsid protein, but rather had a profound effect upon virion assembly and genomic encapsidation. The connection between viral genome dimerization and packaging has been previously described for retroviruses such as in Human Immunodeficiency Virus 1 (Sakuragi, Ueda et al. 2003, Russell, Liang et al. 2004, Lu, Heng et al. 2011) and murine leukemia virus (D’Souza and Summers 2004, Hibbert Catherine, Mirro et al. 2004), as well as for multi-segmented dsRNA orbiviruses such as bluetongue virus (Roy 2017) and rotavirus (McDonald and Patton 2011). For FHV and other alphanodaviruses, we speculate that the RNA1-RNA2 heterodimerization contributes to virus packaging efficiency via non-covalent coupling of each RNA that subsequently ensures stoichiometric RNA encapsidation.

One unexpected finding was that HD2 mutant produced significantly more sgRNA3 than wt FHV. It has previously been demonstrated that two cis-acting elements on RNA1 (nt.1229-1239 and nt.2282-2777) regulate the production of sgRNA3 (Lindenbach, Sgro et al. 2002), whose replication further trans-activates the replication of RNA2 (Albariño, Eckerle et al. 2003, Eckerle, Albarino et al. 2003). The RNA3-dependent replication of RNA2 reciprocally suppresses RNA3 synthesis via an unknown mechanism (Eckerle and Ball 2002, Eckerle, Albarino et al. 2003). In this study, the discovered heterodimer stem is not adjacent to any known *cis-* or *trans-*acting elements. Nonetheless, the HD2 mutant exhibited increased sgRNA3 and reduced RNA2 replication. This suggests that the region forming the heterodimer stem (RNA1:1094-1102, RNA2:277-285) may also facilitate the RNA3-RNA2 trans-activation or regulation of RNA3 expression and may have further roles in maintaining intracellular RNA1-RNA2 stoichiometry.

It is interesting that, although HD1 was designed to contain more disruptive mutations to the heterodimer interaction site, HD2 exhibited more disruptive phenotypes, such as decreased virion yield (**Figure 5c**), perturbed RNA1:RNA2 encapsidation ratios (**Figure 5h**) and reduced thermostability (**Figure 6e**). This suggests that the genotypic introduction of single-stranded bases does not guarantee base-pair disruptions in virus, as the introduced single stranded RNA bases can be base-paired elsewhere. This is also supported by our DMS-MaPseq results (**Figure 4b**).

### XL-ClickSeq and dsRNA probing

NGS-coupled RNA structure methods such as SHAPE-MaP (Smola and Weeks 2018) and DMS-MaPseq (Zubradt, Gupta et al. 2017) have enabled high-throughput RNA secondary structure mapping on genomic and transcriptomic scales, in both *in vitro* and *in vivo* settings. Their correspondent chemicals (SHAPE reagents such as 1-methyl-7-nitro-isatoic anhydride (1M7) and dimethyl sulfide) effectively methylate single-stranded RNA bases that can subsequently be detected by NGS analyses (Wells, Hughes et al. 2000, Weeks and Mauger 2011). As such, double-stranded RNA is typically inferred reciprocally, with the support of computational thermodynamic prediction algorithms (Zhou and Routh 2020). Psoralen-based crosslinking methods thus provide an alternative and direct approach to specifically probe dsRNAs. Here, we developed a novel method (XL-ClickSeq) as an alternative to ssRNA chemical probing methods (such as SHAPE-MaP and DMS-MaPseq) and dsRNA crosslinking and proximal ligation methods (Lu, Gong et al. 2018, Ziv, Gabryelska et al. 2018). XL-ClickSeq specifically probes crosslinked dsRNA regions with a highly efficient protocol that only requires a single crosslinking step prior to routine to NGS library constructions using ‘ClickSeq’. XL-ClickSeq reveals dsRNA sites by marking the approximate location of the predicted crosslinks, therefore providing a near-nucleotide resolution of the covalent bond(s) that resides in dsRNAs (we estimate within 10nts. of the XL-ClickSeq site). Of note, XL-ClickSeq does not provide the sequence of the crosslinked counterpart. This is in contrast to approaches such as PARIS (Lu, Gong et al. 2018) and COMRADES (Ziv, Gabryelska et al. 2018) that sequence both strands of a crosslinked dsRNA after proximity ligation, allowing for identification of long-distance and inter-molecule RNA-RNA interactions. Hence XL-ClickSeq still relies on thermodynamic prediction to discover potential long-distance RNA-RNA interactions, as demonstrated in this study.

### High throughput virion thermostability assay

In this study, we presented a high-throughput method to track the change in permeability of virions to intercalating dyes during thermo-denaturation. The approach requires a fluorescent nucleic acid stain with affinity to both ss- and ds-RNA (such as ‘GelStar’ in this study) and can be conducted in any fluorescence-based qRT-PCR machine with correspondent emission/excitation filters. As such, this approach could be easily applied in wide range of viruses. There are two essential pre-requisites to ensure the accurate interpretation of virus thermostability: (1) purity of viruses must be maximized as any co-purified or non-encapsidated nucleic acid can yield a high background fluorescence. In this study, each virus particle was subjected to five rounds of purification (1X PEG precipitation, 2X sucrose gradient ultracentrifugation and 2X molecular weight filtration), as well as two rounds of RNase/DNase digestion. (2) After each heating step, it is important to measure fluorescence at a constant temperature (37°C in this study) to avoid temperature-induced absorbance change of the nucleic acid stain. With this simple high-throughput method, we discovered that the mutant viruses with disrupted heterodimer stem exhibited reduced thermostability (**Figure 6b**). This is consistent with findings from molecular assays (**Figure 6a**).

It has been speculated that FHV RNA secondary structures may serve as scaffold to maintain structural stability (Fisher and Johnson 1993). We have also previously demonstrated using NGS-based RNA-protein cross-linking approaches that mature FHV virions contain extensive RNA-capsid interaction sites and that these sites are enriched in double-stranded regions of the RNA genome (Zhou and Routh 2020). The reduced thermostability of FHV mutants demonstrates that disruption of RNA secondary structure of encapsidated viral genome has a direct impact on the virus supramolecular structure and particle thermostability. As there are likely multiple heterodimer sites in the encapsidated RNAs, it is foreseeable that these RNA1-RNA2 interactions collectively contribute to the overall structure stability and physical integrity of FHV virions.

## Conflict of Interest Statement

A.R. is a co-founder and owner of ‘ClickSeq Technologies LLC’, a Texas-based Next-Generation Sequencing provider offering ClickSeq protocols and downstream analyses such as those presented in this manuscript.

## Funding Acknowledgements

This work was funded by: NIH grant R21AI151725 to ALR and National Institute of Allergy and Infectious Diseases U54AI150472 subcontract to ALR.

## Data Availability

The raw sequencing data of this study are available in the NCBI sequence read archive (SRA) with accession number: PRJNA858427.

## Methods

### Cell culture and virus

Wild-type (wt) Flock House virus (FHV) was generated from transfecting *Drosophila melanogaster* (S2) cells. The detailed protocol regarding transfection, S2 culture maintenance and virus passaging was described previously (Zhou and Routh 2020). In most of this study, wt virus and mutants were purified with standard methods (Zhou and Routh 2020), which sequentially consists of 4% polyethylene glycol (PEG) 8000 precipitation, RNase/DNase digestion, sucrose gradient ultracentrifugation and 100K MWCO polyethersulfone (PES) membrane filtration. During packaging specificity and thermostability experiments, quintuple-purified virus stocks were used to ensure minimum non-viral content: following standard purification, the purified viruses further underwent additional RNase/DNase digestion, sucrose gradient purification, and PES membrane filtration.

### Heterodimer formation

Purified FHV virions (in 50 mM HEPES) were heated to 65°C for 10 min in a thermocycler. This was followed by slow cooling at a pace of -2 °C per 5 sec until 4 °C. A number of RNA extraction methods were experimented (see **Discussions**). In this study, all heterodimer RNAs were extracted and retained with Quick-RNA Viral kit (Zymo Research) with standard manufacturer’s protocol. The extracted heterodimer was kept at 4 °C until further analyses. The electrophoresis of heterodimer RNA was conducted under non-denaturing conditions, with 1% agarose, 1 × lithium acetate borate buffer and a custom loading dye consisting of 2.5% Ficoll-400, 6.6 mM Tris-HCl ph 8.0, 0.01% OrangeG.

### Northern blot

The heterodimer bands were cut from agarose gel (**Figure 4a**) and the RNA content was recovered with Zymoclean Gel RNA Recovery Kit (Zymo Research) using standard procedures. Equal amounts of RNA were heat denatured (65°C 10 min). Two fluorescently labelled ssDNA probes were designed to specifically target the conserved FHV +RNA1/+RNA3 and +RNA2 sequences, respectively (RNA1/RNA3: Cy5-GAGTGTTGGTTTTGCCTCCT; RNA2: Cy3-GAAACGCCAAACCAGGTTGACTTAATCTGGTTAGCGCCGCCATGTTCAT). Electrophoresis, membrane transfer and hybridization were conducted with NorthernMax (Ambion) kit and protocol with modifications: (1) during electrophoresis, a dye-less loading buffer was made to prevent fluorescent interference; (2) transferred nylon membrane was UV crosslinked with UV stratalinker 2400 (1200 μJoules); (3) hybridization was conducted with ULTRAhyb-Oligo buffer instead of ULTRAhyb at 42°C overnight, with 4 μL of each probe at 10 mM; (4) post-hybridization, membrane was sequentially washed twice with low stringency wash (5 min RT), once with low stringency wash (2 min 42°C), twice with TBST (tris-buffered saline and 0.1% Tween-20) buffer wash (5 min RT) and twice with TBS (tris-buffered saline) buffer (5min RT); (5) membrane was exposed on Typhoon (FLA9500) to detect fluorescence. Densitometry was conducted with ImageJ (NIH). Statistical assays were conducted with one tailed paired t-Test for means, with α = 0.05, N ≥ 3 biological replicates.

### Oxford Nanopore Technology (ONT) MinION sequencing

The RNA extracted from defective FHV particles was sequenced with ONT MinION. RNA was reverse transcribed, PCR amplified and ligated with KIT SQK-LSK109 with standard methods and sequenced with a MinION R9.4 flowcell. The detailed library protocol, software settings, data processing and genome alignment methods are previously described (Jaworski and Routh 2017).

### AMT crosslinking

FHV RNAs extracted from heated particles were crosslinked with 4′-aminomethyl trioxsalen hydrochloride (AMT). 1 μg of purified RNA is mixed with AMT (at 20 μg/mL) and placed 5 cm underneath a 365 nm UV light (3UV-38, UVP) for 38 min on ice in a dark room. This yielded approximately 0.15J/cm^2^ of energy. The crosslinked RNA was purified with RNA Clean & Concentrator (Zymo Research) to remove excessive AMT.

### XL-ClickSeq

AMT-crosslinked RNA was directly used as template during reverse transcription. Standard ClickSeq protocol (Jaworski and Routh 2018) was carried out. In brief, 250 ng of templated RNA was supplemented with a mixture of azido-NTP:dNTP (1:5 molecular ratio) during random primed reverse transcription. This is followed by “Click-ligation” (Kolb, Finn et al. 2001) with an 5’-hexynyl-functionalized Illumina adapter (IDT). The adapter ligated cDNA fragments were subsequently PCR amplified to complete Illumina adapter sequences and to incorporate samples indexes/barcodes. The ClickSeq libraries of non-crosslinked FHV RNAs were constructed the same way as for the control.

### Bioinformatics

The bioinformatics of XL-ClickSeq followed a published bioinformatic pipeline (Zhou and Routh 2020). In brief: the raw Illumina sequencing data were sequentially prepared by trimming Illumina adapter (*cutadapt* (Martin 2011): -b AGATCGGAAGAGC -m 40), trimming of nucleotides adjacent to the triazole-linkage in the ‘click-linked’ cDNA using*FASTX toolkit* (http://hannonlab.cshl.edu/fastxtoolkit/index.html: (fastx trimmer -Q33 -f 7), and filtering read quality (fastq_quality_filter -Q33 -q 20 -p 96). End-to-end alignment to the FHV genome (NC 004146, and NC 004144 for RNA1 and RNA2, respectively) was conducted (*Bowtie* (Langmead, Trapnell et al. 2009): -v 2 --best). The genome coverage data were extracted from the aligned reads (SAMtools (Li, Handsaker et al. 2009): view/sort/mpileup), and the genome coverage of crosslinked heterodimer was compared with non-crosslinked RNA.

For both DMS-MaPseq and HD mutant packaged RNA characterization, the generated azido-tagged cDNAs were “click”-ligated with an Illumina adapter with 12 random nucleotides to act as Unique Molecular Identifiers for de-duplication and control of of PCR bias. Raw reads were processed to remove Illumina adapter and to assign unique identifier (*fastp* (Chen, Zhou et al. 2018): -a AGATCGGAAGAGC -U --umi_loc read1 --umi_len 14 --umi_prefix umi -l 30). For DMS-MaPseq, *Bowtie2* (Langmead and Salzberg 2012) was used to align reads to FHV genome to allow gapped alignment (--local). This is followed by de-duplication to remove PCR bias (*umi-tools* (Smith, Heger et al. 2017): dedup --method=unique) and extraction of mutation rate with a minimum nucleotide coverage of 1000 (SAMtools (Li, Handsaker et al. 2009): view/sort/mpileup). For packaged RNAs of mutant viruses, *Hisat2* (Kim, Paggi et al. 2019) was used for alignment with default settings to FHV genome first, while the unmapped reads were further aligned to *Drosophila melanogaster* genome (release 6 (Hoskins, Carlson et al. 2015), GCA_000001215.4). This is followed by de-duplication and read extraction as stated above. To characterize the packaged RNAs of viruses, statistical assays were conducted with one-tailed Student t-Test (equal variance), with α = 0.05, N ≥ 3 biological replicates.

### RNA structure thermodynamic prediction

Free energy based thermodynamic prediction was conducted with *RNAstructure Web Server* (Reuter and Mathews 2010), with “bifold” algorithm to allow both inter- and intra-molecular base pairs. 7 RNA1 heterodimer candidate sites (whose flanking sequences composed into 5 candidate regions) and 6 RNA2 candidate sites (whose flanking sequences composed into 6 candidate regions) were cross matched for RNA secondary structure prediction (**Figure S3**). The predicted structure file was re-organized, re-colored for graphical purposes with *StructureEditor*, which is provided by *RNAstructure* suite.

### Mutant virus cloning, transfection and passaging

The primers used to generate the In-Fusion cloning (TaKaRa) fragments were listed in **Table S1**. Universal upstream (TGCATAATTCTCTTACTGTCATGCCATCCGTAAG) and downstream (TAAGAGAATTATGCAGTGCTGCCATAACCATG) primers were used to target the backbone of pMT plasmids (Invitrogen). Overlapped PCR fragments were generated (Phusion High-Fidelity DNA Polymerase, NEB) and cloned into competent cells with standard In-Fusion HD cloning techniques. The plasmids contained mutated viral sequences were sanger sequenced to confirm the mutation. Equimolar RNA1 and RNA2 plasmids containing mutated heterodimer sequences were used to co-transfect S2 cells with Lipofectamine 3000 (Invitrogen) with standard protocols and induced with copper sulfate 24 h post transfection. The generated P0 viruses were serially passaged up to P2 with 3 days interval.

### DMS-MaPseq

Dimethyl sulfate (DMS) RNA methylation method was described previously (Zhou and Routh 2020). In brief, DMS was used at 5% final concentration with purified viruses for 5 min at 30 °C. This was followed by 5 min quenching with excess volume of 10 mM Tris pH 7.4 and 30% 2-mercaptoethanol (BME). RNA was extracted from DMS-treated virions and RNAseq libraries were constructed using ClickSeq as previously stated (Zhou and Routh 2020). For each virus (wt/HD1/HD2), the nucleotide mutation rate of DMS-treated virus was compared with that of the untreated and corresponding control virus to generate signal. To distinguish DMS-MaPseq signals from background, a background threshold was applied as previously (Zhou and Routh 2020), which is determined as the highest 5% mutation rate of the corresponding control.

### Relative cytotoxicity

In this study, the relative cytotoxicity of each virus was measured by alamarBlue cell viability assay of P0 transfected S2 cells, as described previously (Zhou and Routh 2020). In brief, 25k naïve S2 cells were seeded in black 96-well plate, transfected with 100 ng plasmids of wt or mutant virus genome, and induced with copper sulfate. 2 days post induction, alamarBlue cell viability reagent (ThermoFisher) was supplemented. Cells were incubated for 4 hours, before fluorescence detection with an EnSpire plate reader (PerkinElmer) at 560 nm excitation and 590 nm emission. The relative fluorescence, which indicates viable cell count, then was normalized reverse-ratiometrically to relative cytotoxicity, with mock transfection = 0% and FHV wt transfection = 100%. Statistical assays were conducted with one tailed Student t-Test (equal variances), with α = 0.05, N ≥ 6 biological replicates.

### Western blot

Three days post transfection, equal volume of transfected S2 cells (P0 virus) were used to inoculate naïve cells to generate P1 viruses. 24 h post inoculation, 100 μL culture were collected. Cell and supernatant fractions were separated with centrifugation (1000 × g, 10 min, 4 °C). Cell fraction was washed twice and resuspended into 30 μL with 1 × phosphate-buffered saline (PBS) and 1 × cOmplete proteinase inhibitor (Roche). Supernatant fraction was supplemented with 1 × cOmplete and reduced to 30 μL with vacuum centrifuge. Entire 30 μL of both cellular and supernatant fractions were loaded on to a Bolt 4–12% Bis–Tris Plus Gels (Invitrogen). Details regarding membrane transfer and western blotting were documented previously (Zhou and Routh 2020). In brief, a rabbit anti-FHV polyclonal antibody and an Alexa Fluor 488 goat anti-rabbit IgG (Invitrogen) were used to probe FHV alpha, beta, and gamma peptides. Prior to membrane transfer, part of SDS-PAGE gel was cut and stained (Coomassie brilliant blue R-250) to highlight α-tubulin (55 kDa) as a loading control. Labelled membrane was visualized with Typhon (FLA9500) and densitometry was conducted with ImageJ (NIH).

Statistical assays were conducted with one tailed Student t-Test (equal variances), with α = 0.05, N ≥ 4 biological replicates.

### High-throughput thermostability assay

A high-throughput virion thermostability assay was developed by repurposing a qRT-PCR program. In this assay, it is important to ensure the purity of virions and the minimum of co-purified, non-encapsidated cellular RNA. As detailed above, all viruses were quintuple purified which consists of multiple purification as well as two rounds of RNase A/DNase I digestion. A 50 μL mixture consisting of 50 μg of purified particle, 50 mM HEPES (pH 7.2), 1 × nucleic acid stain (GelStar, Lonza) was added to a 96 well qRT-PCR plate. A thermocycling program was set on a qRT-PCR machine (Roche LightCycler 480II) (**Figure 6b**), which comprises of a 1 min incubation cycle with gradually increased temperature (1 °C per cycle, from 37 °C – 95 °C), and a 10 s cooling at 37 °C. Fluorescence reading was taken at the end of each cooling to avoid any temperature induced absorbance change of nucleic acid dye. The observed temperature-to-fluorescence curve of different particles can be transformed to a fluorescence-to-temperature formulas, which is generated by converting the polynomial (order = 2) trendline of average fluorescence change of (from 47 °C – 62 °C). The melting temperature of each virus particle (Tm, which is defined as the temperature to reach 50% fluorescence increase) can be calculated for each replicate thereafter. Statistical assays were conducted with one tailed Student t-Test (equal variances), with α = 0.05, N = 12 (each sample consisted of 2 biological replicates, each biological replicate consisted of 6 technical repeats).

## Notes

https://www.ncbi.nlm.nih.gov/sra/?term=PRJNA858427

