## Supplementary material for "Bipartite viral RNA genome heterodimerization influences genome packaging and virion thermostability": Suppl.Figures

### Supplementary Data

**Supplemental Table 1. Primers used for HD mutants**

| | | | |
|---|---|---|---|
| HD1 | RNA1 | Forward primer | AGAGCGTGAACGCTAGGCTTATCGGTATGGGA |
| | | Reverse primer | TAGCGTTCACGCTCTGGGTGGCGGATAATC |
| | RNA2 | Forward primer | AGATAGATTTGAAGGCAAAGTGGTCAGCCGAAAG |
| | | Reverse primer | CCTTCAAATCTATCTGGTATTCCCTTACCGGGGT |
| HD2 | RNA1 | Forward primer | AGTCTGTGAACGCTAGGCTTATCGGTATGGGA |
| | | Reverse primer | TAGCGTTCACAGACTGGGTGGCGGATAATC |
| | RNA2 | Forward primer | AGATCGTTTTGAAGGCAAAGTGGTCAGCCGA |
| | | Reverse primer | CCTTCAAAACGATCTGGTATTCCCTTACCGGGGTC |

**Supplemental Table S1. Primers designed for In-Fusion Cloning (TaKaRa) to create mutants HD1 and HD2.** Red bases represents mutated nucleotides.

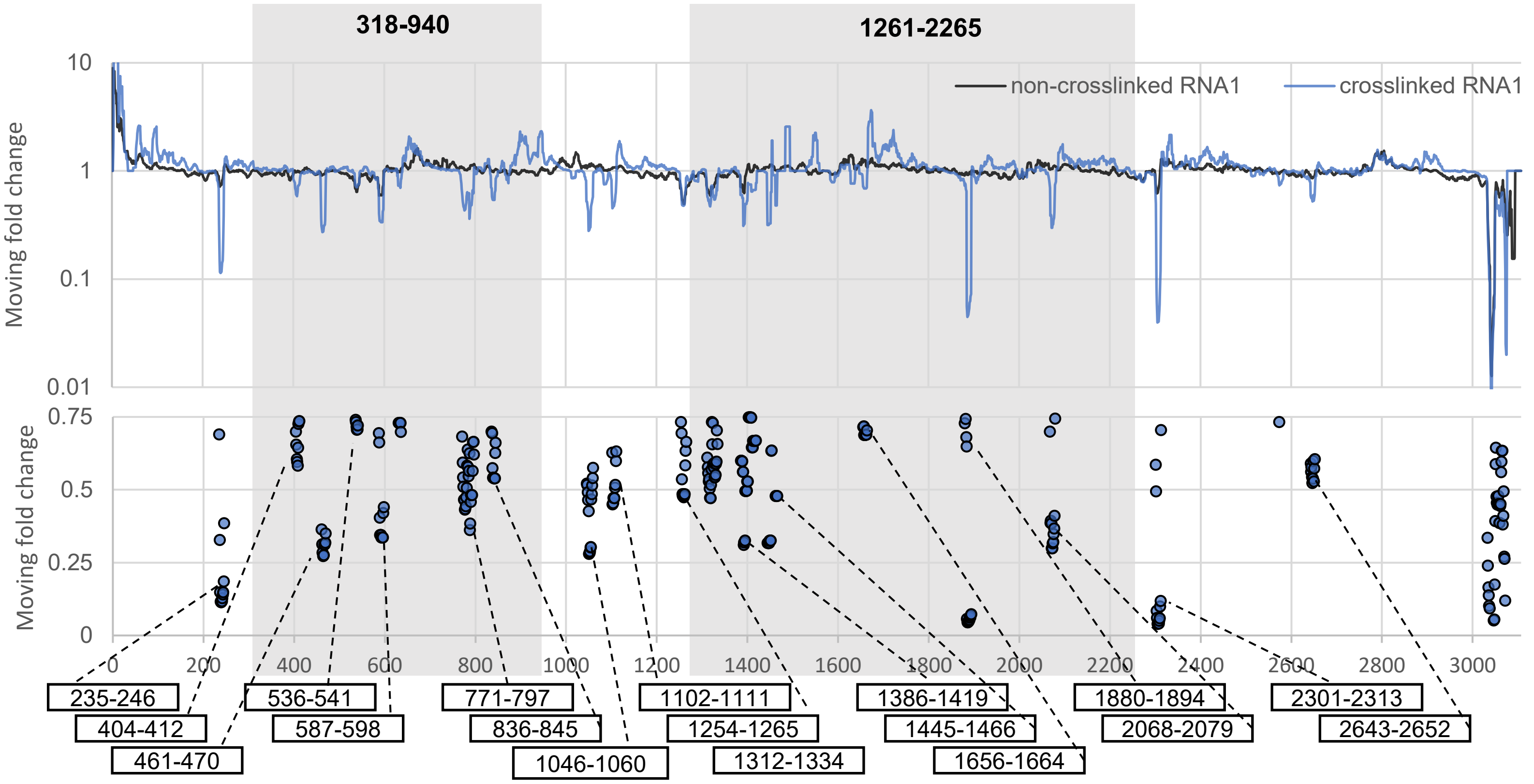
318-940
1261-2265
non-crosslinked RNA1
crosslinked RNA1
Moving fold change
10
1
0.1
0.01
Moving fold change
0.75
0.5
0.25
0
0
200
400
600
800
1000
1200
1400
1600
1800
2000
2200
2400
2600
2800
3000
235-246
404-412
461-470
536-541
587-598
771-797
836-845
1046-1060
1102-1111
1254-1265
1312-1334
1386-1419
1445-1466
1656-1664
1880-1894
2068-2079
2301-2313
2643-2652

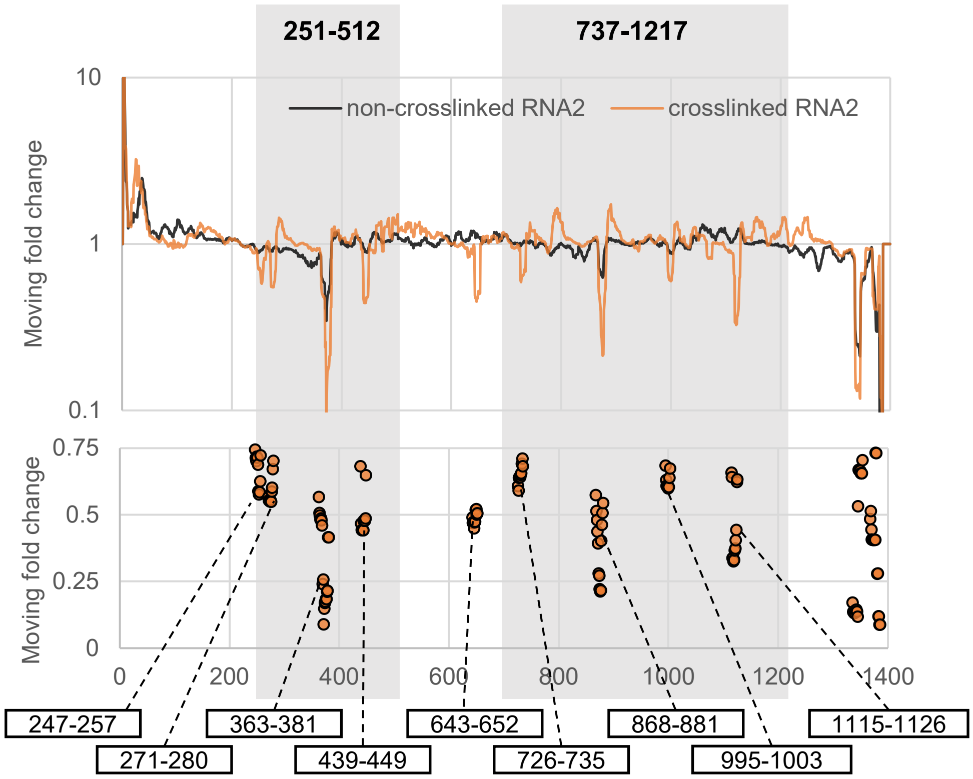
251-512
737-1217
non-crosslinked RNA2
crosslinked RNA2
Moving fold change
10
1
0.1
Moving fold change
0.75
0.5
0.25
0
0
200
400
600
800
1000
1200
1400
247-257
271-280
363-381
439-449
643-652
726-735
868-881
995-1003
1115-1126

**Supplemental Figure S1. Potential heterodimer sites.** After XL-ClickSeq, the coverage differences between crosslinked heterodimer and non-crosslinked control are shown as moving fold changes with a 10 nts. Interval. The crosslinked dsRNA regions are clearly indicated by loss-of-coverage. Consecutive nucleotides (greater than 6) that have a greater than 25% moving fold change are shown in the scatter plots. For both FHV RNA1 and RNA2, two major deletions can be found on defective genomes (RNA1:318-940, 1261-2265; RNA2: 251-512, 737-1217, **Supplemental Figure S2**). As FHV heterodimer is formed with D-RNA1 but not D-RNA2 (**Figure S3**), it is possible to narrow the selection of candidate heterodimer sites. In this study, RNA1 heterodimer sites were selected outside the deleted regions, and RNA2 heterodimer sites were selected inside the deleted regions.

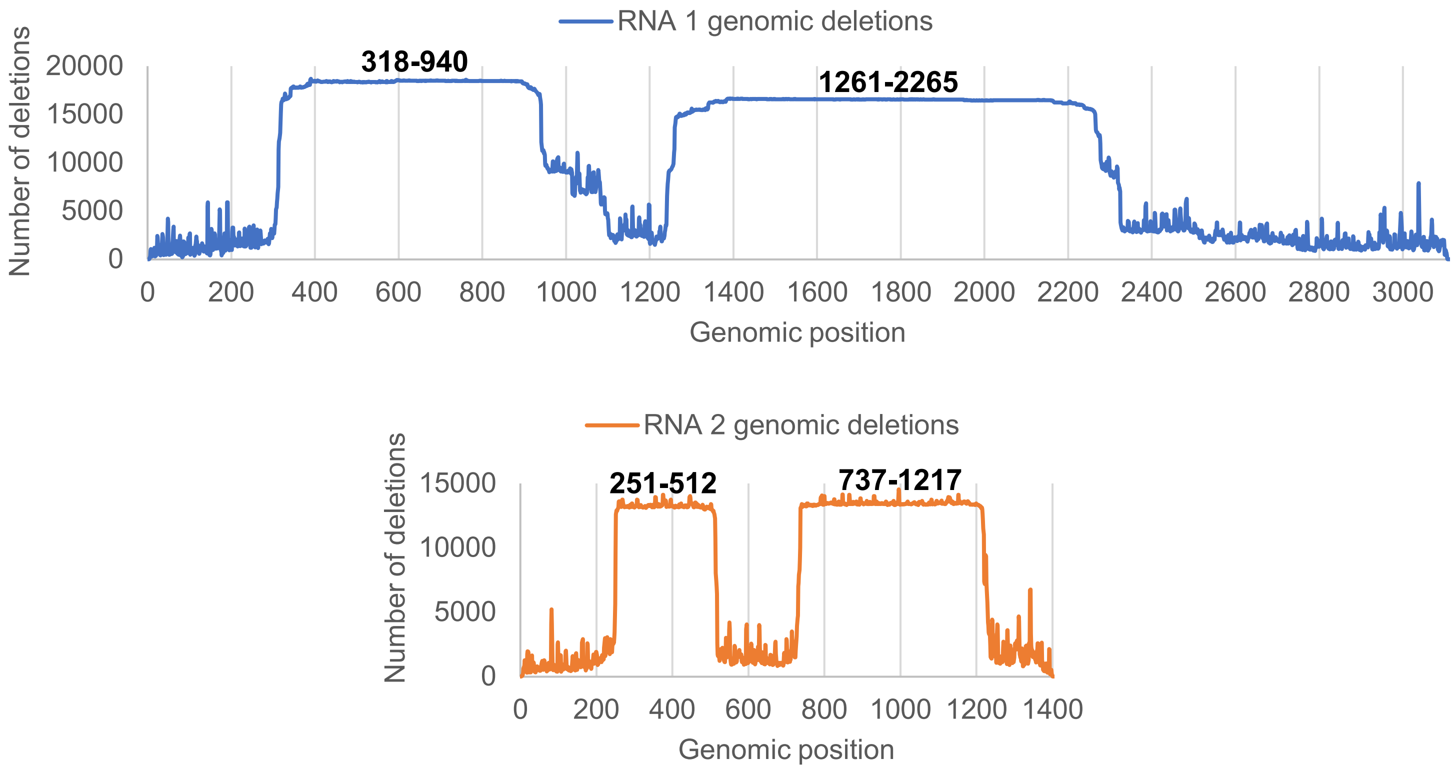


**Supplemental Figure S2. FHV P7 defective RNA genome is characterized by major deletions.** The RNAs were extracted from P7 FHV DI particles and subjected to nanopore sequencing to reveal genomic deletions. P7 FHV features significant deletions events at nt. 318-940 and nt. 1261-2265 on RNA1 and nt. 251-512 and nt. 717-1217 on RNA2.

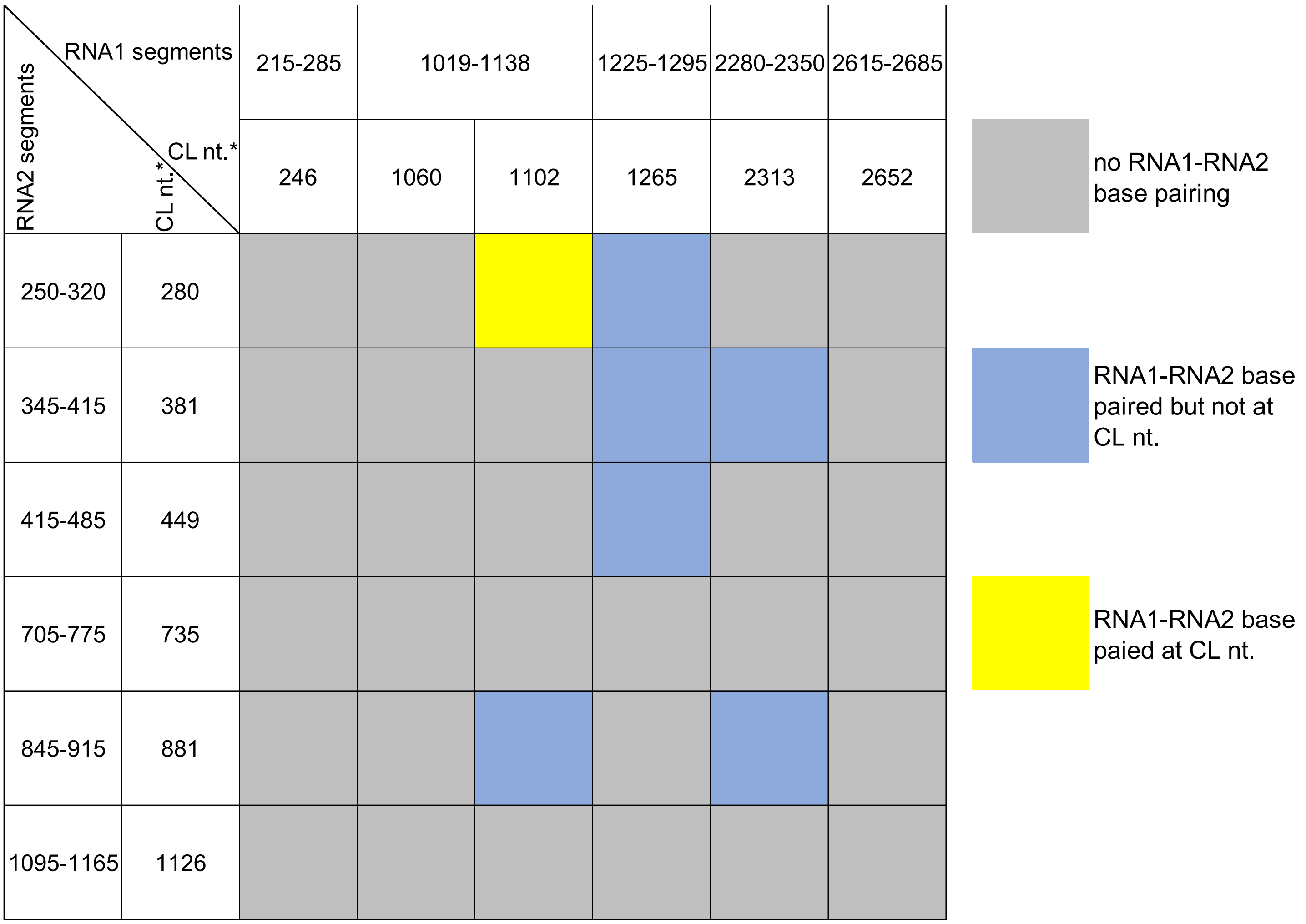


**Supplemental Figure S3. Inter-RNA base pairing prediction of candidate heterodimer sites.** The candidate RNA1 and RNA2 heterodimer sites, including their flanking sequences (71-120nts.), were used to predict intermolecular thermodynamic base pairing potentials with RNAstructure (REF). Each RNA1 segment was cross-matched with every RNA2 segments to evaluate whether substantial inter-RNA base-pairing can form and whether the base-pairing is located near the predicted crosslink (CL) sites. Most RNA segments of candidate site were either unable to form intermolecular base pairs (grey cells) or formed base pairs too distant (>20 nts.) from the predicted candidate CL sites (blue cells). Only RNA1:1019-1138 and RNA2:250-320 formed multiple intermolecular stems at the proximity of predicted CL sites (RNA1:1102, RNA2:280), which also contains inter-stranded pyrimidines available for AMT crosslinking (yellow cell, structure illustrated in **Figure 4a**). *: CL nt. only indicates the approximate crosslinking site, as the precise location of covalent crosslinking bond can reside in proximity of this site.

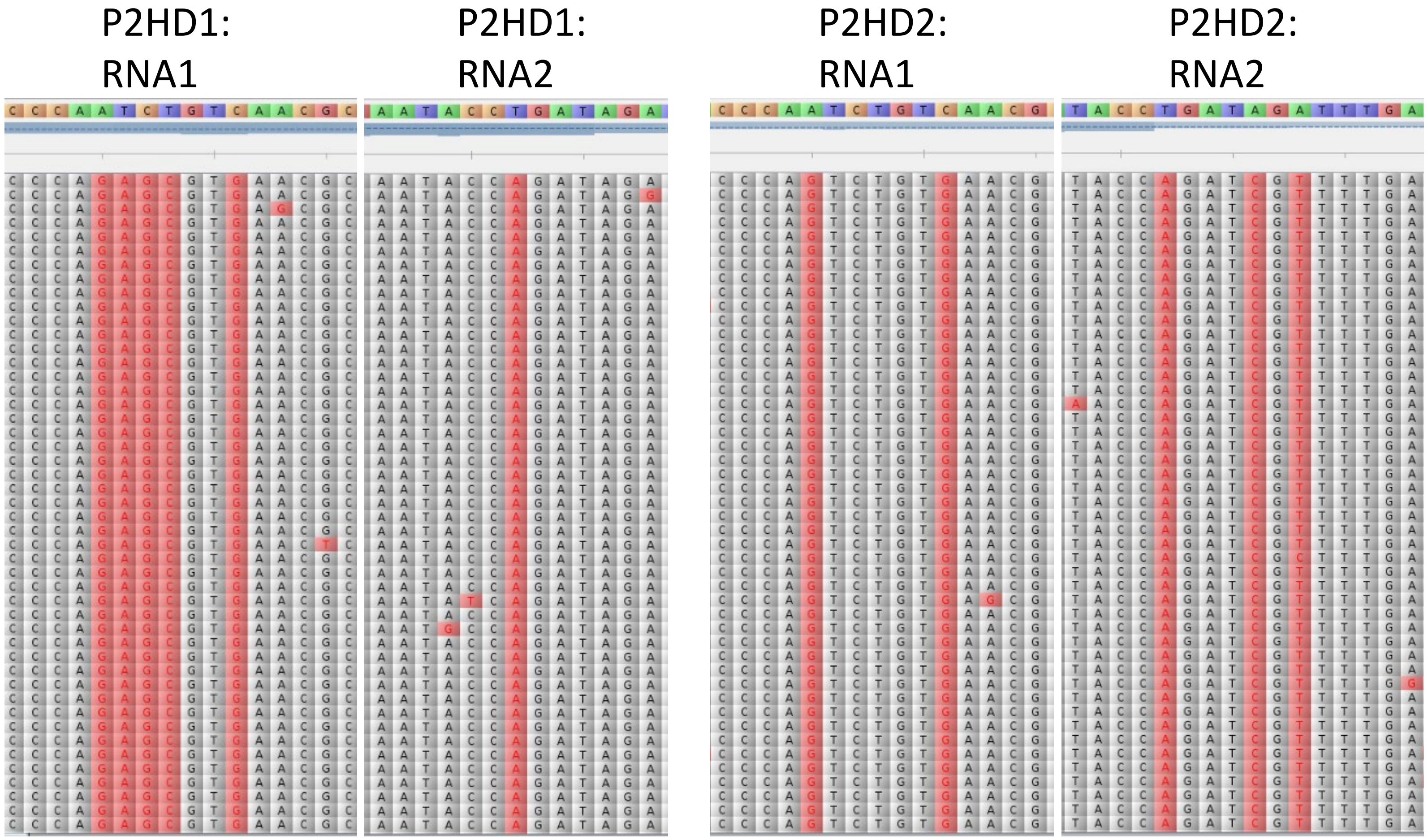


**Supplemental Figure S4. Mutations in HD1 and HD2 are retained up to P2 viruses.** Passage 2 mutants (P2HD1 and P2HD2) were purified and sequenced. Reads of mutants were mapped to the correspondent genomes, and nucleotide variants were visualized in Tablet (REF) by comparing with wtFHV genome. Mutated nucleotides of HD1 and HD2 are highlighted in red and well reserved in P2 viruses. Further passages are not pursued in this research.

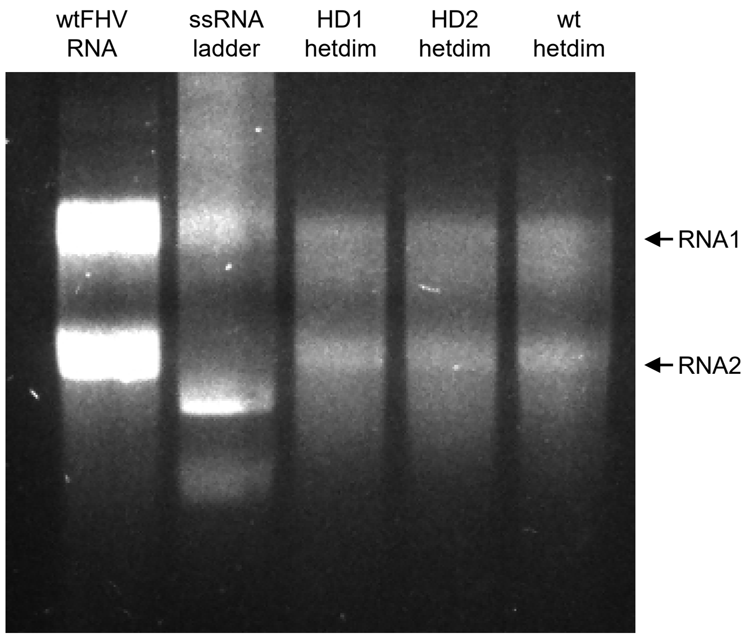


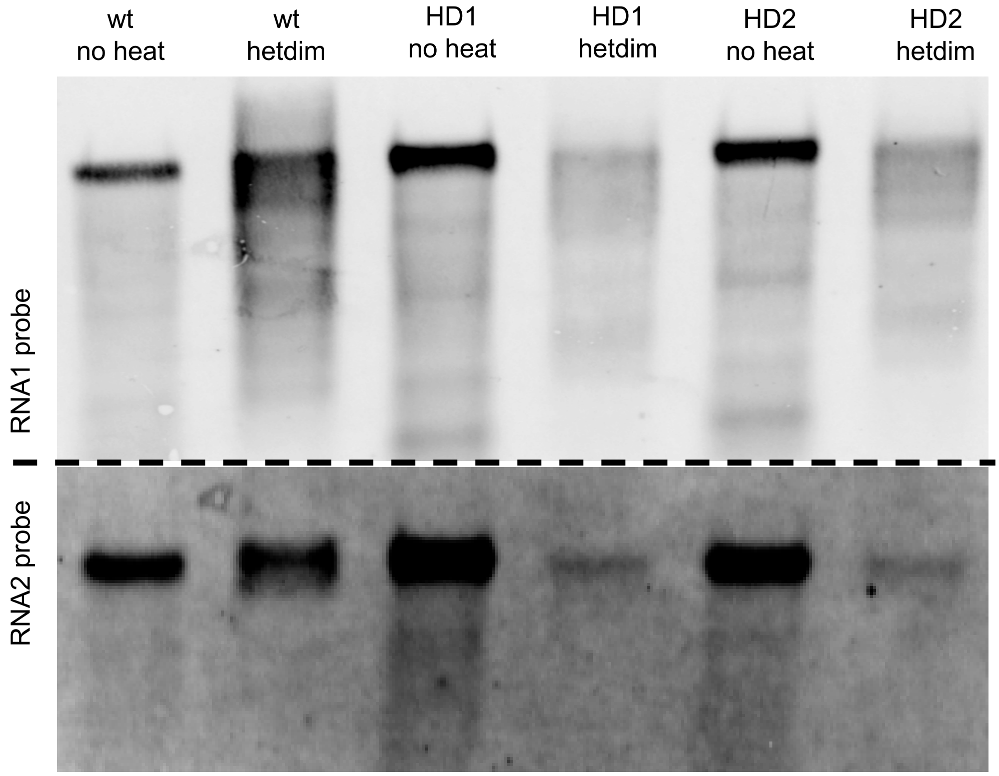


**Supplemental Figure S5. HD1 and HD2 mutants heterodimers consists of both RNAs.** Heterodimer bands were cut from HD1, HD2 and wtFHV (**Figure 4c**). RNAs were recovered from native agarose gel and subjected to heat denaturing. The denatured heterodimers were then run on a native agarose gel **(a)** and denaturing gel for northern blot **(b)**. The presence of both RNA1 and RNA2 can be detected from the denatured heterodimer (hetdim) band of both HD1 and HD2.

a

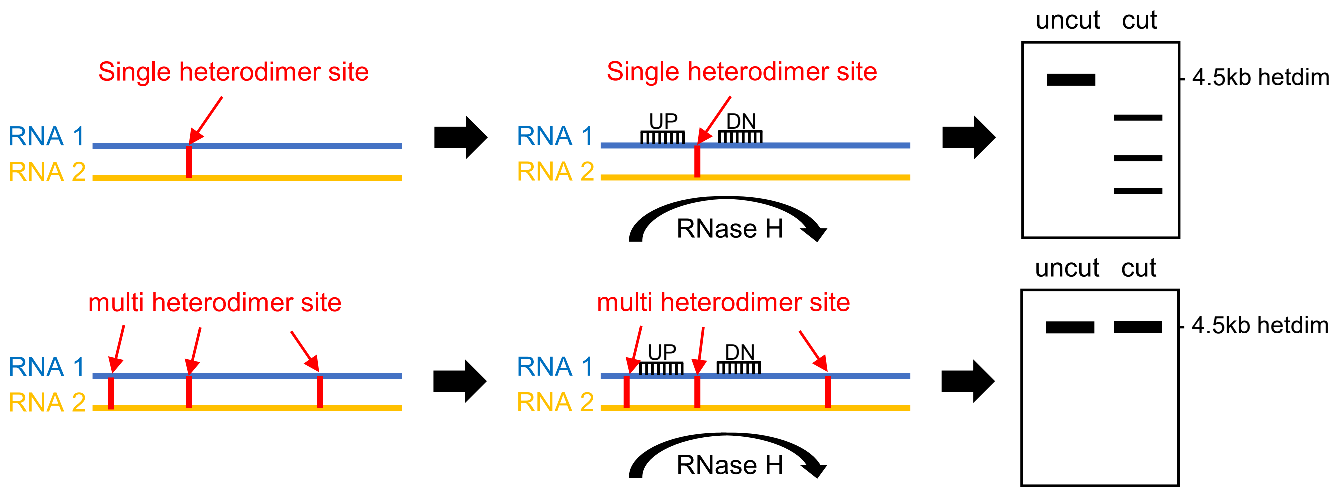
Single heterodimer site
RNA 1
RNA 2
Single heterodimer site
UP
DN
RNA 1
RNA 2
RNase H
uncut cut
4.5kb hetdim
multi heterodimer site
RNA 1
RNA 2
multi heterodimer site
UP
DN
RNA 1
RNA 2
RNase H
uncut cut
4.5kb hetdim

b

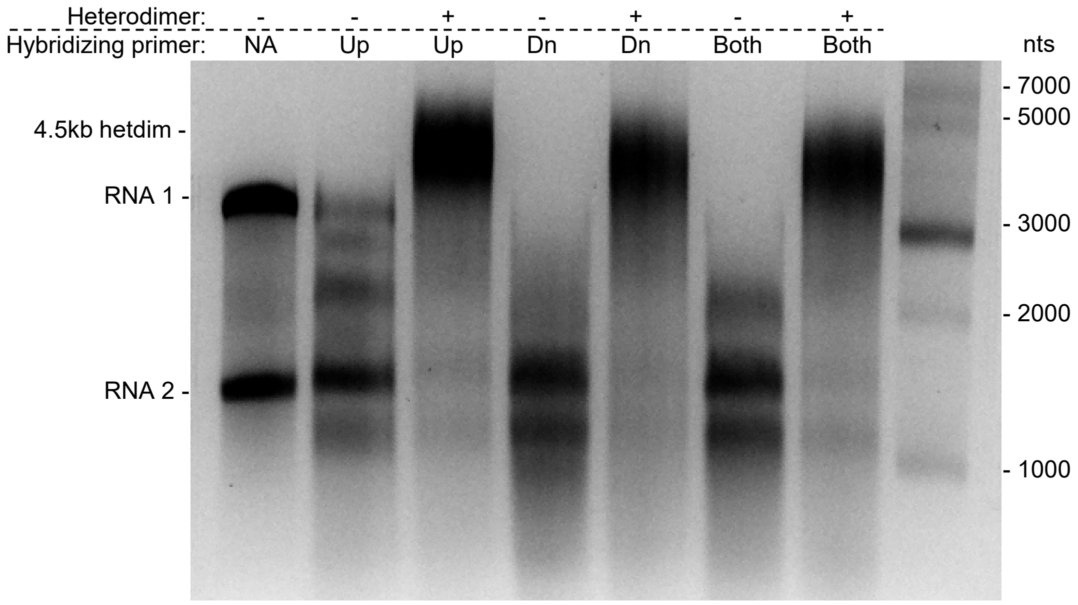
Heterodimer: - - + - + - +
Hybridizing primer: NA Up Up Dn Dn Both Both
nts
4.5kb hetdim -
RNA 1 -
RNA 2 -
- 7000
- 5000
- 3000
- 2000
- 1000

**Supplemental Figure S6. RNase H digestion confirmed multiple heterodimer sites beyond the predicted stem. (a)** the scheme of RNase H molecular assay to determine if FHV heterodimer contains multiple RNA1-RNA2 interactions. If there is only one heterodimerizing interaction, heterodimer can be fully disassociated into small molecular RNAs after RNase H digestion at both up- and down- stream of the heterodimer site. If there are multiple heterodimer sites, RNase H digestion at a single site will not fully disassociate heterodimer.**(b)**Two hybridizing DNA oligos were designed to bind RNA1 at up- or down- stream of the predicted heterodimer stem (Up: 5'-AATCTGAGCATGCTCCCCTT-3'; Dn: 5'-AATCATGGATGTGTATTGCG-3'). 500 ng of RNA was hybridized with 5µM oligomer(s) with 50 mM HEPES (pH 7.2) after heating (65°C 10 min) and slow cooling. Hybridized RNA was subjected to RNase H digestion (5U, 37°C 10min) and analyzed on a non-denaturing agarose gel. Crosslinked heterodimer retained duplex formation after digestion, with up-, down- stream oligomers or both. This indicates that there are other heterodimer sites in addition to the 9 bps stem predicted in this study.

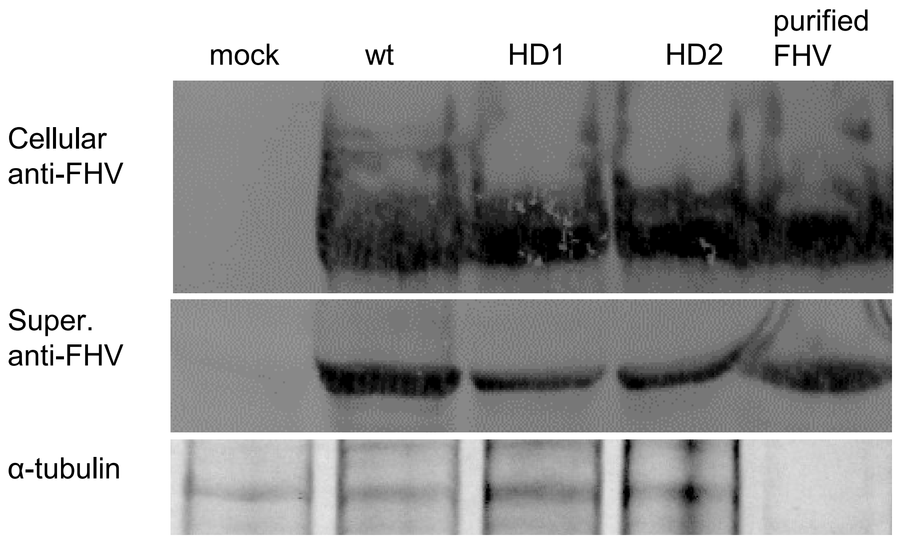


**Supplemental Figure S7.** Western blot revealed differential capsid accumulations from cell or supernatant fractions. Two days post infection of P1 viruses, 50uL of cell/supernatant mixture was collected and cellular fraction was separated from supernatant fraction by centrifugation and PBS washes. The correspondent fractions were run on SDS-PAGE and western blotted with polyclonal FHV antibody to highlight capsid yields. The SDS-PAGE section contains alpha-tubulin (55 kDa) was cut and Coomassie blue stained as a loading control.
